# Reconstitution of ORP-mediated lipid exchange process coupled to PI(4)P metabolism

**DOI:** 10.1101/2023.08.04.551917

**Authors:** Nicolas Fuggetta, Nicola Rigolli, Maud Magdeleine, Agnese Seminara, Guillaume Drin

## Abstract

Lipid distribution in the eukaryotic cells depends on tight couplings between lipid transfer and lipid metabolism. Yet these couplings remain poorly described. Notably, it is unclear to what extent lipid exchangers of the OSBP-related proteins (ORPs) family, coupled to PI(4)P metabolism, contribute to the formation of sterol and phosphatidylserine gradient between the endoplasmic reticulum (ER) and other cell regions. To address this question, we have examined *in vitro* the activity of Osh4p, a representative ORP, between Golgi mimetic membranes in which PI(4)P is produced by a PI 4-kinase and ER mimetic membranes in which PI(4)P is hydrolyzed by the phosphatase Sac1p. Using quantitative, real-time assays, we demonstrate that Osh4p creates a sterol gradient between the two membranes by sterol/PI(4)P exchange as soon as a PI(4)P gradient is generated at this interface following ATP addition, and define how much PI(4)P must be synthesized for this process. Then, using a kinetic model supported by our *in vitro* data, we estimate to what extent PI(4)P metabolism can drive lipid transfer in cells. Finally, we show that Sec14p, by transferring phosphatidylinositol between membranes, can support the synthesis of PI(4)P and the creation of a sterol gradient by Osh4p. These results indicate to what extent ORPs, under the control of PI(4)P metabolism, can distribute lipids in the cell.

## Introduction

Lipids are precisely distributed among the organelles and the plasma membrane (PM) of eukaryotic cells, which is key for the cellular architecture and to support functions such as vesicular trafficking or signal transduction (1, 2). This distribution relies on lipid transfer proteins (LTPs) that carry lipids across the cytosol between cellular membranes in which lipids are synthesized or modified by various enzymes. OSBP and Oxysterol-binding protein (OSBP)-related proteins (ORPs) in human, (called oxysterol-binding homology (Osh) proteins in yeast), represent a family of LTPs involved in the intracellular distribution of key lipids, sterols (cholesterol in human, ergosterol in yeast) and phosphatidylserine (PS), which account respectively for 12-20 % and 2-10% of cell lipids (3–5). These lipids are produced in the ER, but are essentially found in the *trans*-Golgi and the PM (6–12).Sterol guarantees the rigidity and inner organization of these membranes (13, 14) whereas PS, which is a negatively charged phospholipid, contributes to recruiting signaling proteins on the inner leaflet of the PM *via* electrostatic interactions (11, 15, 16). This asymmetrical distribution of sterol and PS can be established by ORPs coupled to the metabolism of a third lipid called PI(4)P but to a degree that remains ill-defined (17).

ORPs have diverse molecular architectures, but all comprise an OSBP-related domain (ORD) with a lipid-binding pocket (18). As first identified with Osh4p and OSBP (19, 20), some of these can host a molecule of sterol and PI(4)P alternately and thereby exchange these lipids between membranes (18). Others, like Osh6p/Osh7p and their closest human homologue ORP5 and ORP8, act as PS/PI(4)P exchangers (21, 22). PI(4)P is mostly synthesized in the *trans*-Golgi and plasma membrane by PI 4-kinases through the phosphorylation of phosphatidylinositol (PI), while in the ER, PI(4)P is hydrolyzed into PI by the phosphatase Sac1. Therefore, a general model is that ORPs extract sterol or PS from the ER, exchange these lipids with PI(4)P at the Golgi or PM, and transfer PI(4)P to the ER ; the maintenance of a PI(4)P concentration gradient at ER/Golgi and ER/PM interfaces by PI(4)P synthesis and hydrolysis fuel continuous exchange cycles, leading to the build-up of sterol and PS in the *trans*-Golgi and/or PM at the expense of the ER, thereby creating asymmetry (17, 18).

Many observations support this model. For instance, *in vitro* studies showed that Osh4p can vectorially transfer sterol between two membranes using a pre-existing PI(4)P gradient (21). In yeast cell, Osh2p supplies the PM with ergosterol to support endocytic processes unless it is unable to trap PI(4)P (23). Osh6p and Osh7p rapidly transfer PS from the ER to the PM unless Sac1p is silenced, which indicates that their activity is driven by a PI(4)P gradient at the ER/PM interface (24). ORP5 and ORP8 display a similar dependency on PI(4)P (22). Likewise, many evidences showed that OSBP delivers sterol to the *trans*-Golgi owing to a PI(4)P gradient at the ER/Golgi interface (20, 25). Finally, data indicate that PI(4)P pools synthesized in endosomes or damaged lysosomes are also used by ORPs to supply these organelles with sterol and PS (26–28). Nevertheless, it has never been fully demonstrated from a biochemical perspective that PI(4)P metabolism can drive lipid exchange cycles via ORPs.

Aside from that, it is unclear whether ORPs perform lipid exchange that are intensive enough to contribute significantly to the asymmetrical distribution in sterol and PS throughout the cell, or only execute limited exchange to fine-tune organelles lipid composition. For instance, in human cells, OSBP can mediate important ER-to-Golgi sterol transfer (20, 25) but it also controls protein recycling at the endosomal level, not by delivering sterol *en masse*, but through a PI(4)P regulation (29). Some clues suggest that, in yeast cells, Osh4p might deliver sterol to the *trans*-Golgi to promote the formation of secretory vesicles(30). However, other data suggest that its main role is to extract PI(4)P from these vesicles to prime them for docking with the PM (31, 32). It is also considered that the role of ORP5 and ORP8 is maybe not to deliver high quantity of PS in the PM but to adjust the PS/PI(4)P balance and PI(4,5)P_2_ levels in this membrane (33, 34).

If the cellular role of ORPs remains difficult to define, this is notably because it is unclear to what extent PI(4)P metabolism can drive their activities. The exchange model posits that one molecule of PI(4)P is consumed for the transfer of one molecule of counterligand. But PI(4)P is in trace amounts in the cells (in yeast, only 0.2% of lipids (35)) compared to sterol and PS, which seems incompatible with the implementation of high lipid exchange. Nevertheless, in yeast lacking Sac1p or all Osh proteins, PI(4)P levels can be multiplied by up to ∼20 (36, 37), which suggests that a pool of PI(4)P, possibly accounting for 2 to 4% of total phospholipids (38) are synthesized by PI 4-kinases and constantly hydrolyzed by Sac1p, to support Osh activities. Also, OSBP has been shown to use half of PI(4)P produced in human cell lines to guarantee 30-60 % of ER-to-Golgi sterol transfer (25). Yet, apart from these few data, we have no quantitative insights into the coupling between ORP-mediated lipid exchange and PI(4)P metabolism.

Because the activity of ORPs depends on PI(4)P, this also raises the question of how the availability of PI is maintained in the cytosolic leaflet of the endosomal, Golgi and plasma membranes (39, 40) to guarantee a constant PI(4)P production. So far, little is known about how PI is exported from the ER, in which it is synthesized (41), to other organelles (39). In yeast, Sec14p supports the production of PI(4)P in the Golgi by the kinase Pik1p and the secretory function of this organelle (35, 42, 43), possibly by transferring PI from the ER to the Golgi membrane *via* PI/phosphatidylcholine(PC) exchange, yet this model is disputed (44, 45). Interestingly, Sac1p and Osh4p are able to directly counteract Sec14p and Pik1p functions (43, 46–48). In human cells, a structurally different class of proteins maintains proper PI(4)P and PI(4,5)P_2_ levels in the Golgi and PM possibly by supplying these membranes with PI (44, 45). Intriguingly, a few clues suggest that some of these PI-transfer proteins support the activity of OSBP at the ER/Golgi interface or between the ER and replication organelles in cells infected by viruses (49, 50). Jointly, these observations suggest that the activity of PI-transfer proteins could be intimately linked to PI(4)P metabolism and ORP-mediated lipid exchange in eukaryotic cells.

To better define the couplings between ORPs activities and PI(4)P metabolism but also PI-transfer processes, we reconstituted an ER/Golgi interface *in vitro* to conduct functional assays in an controlled environment. First, we designed a PI 4-kinase that can be attached to liposomes and efficiently convert PI into PI(4)P in the presence of ATP. Secondly, we measured the ability of Osh4p to perform sterol/PI(4)P exchange between ER-like liposomes functionalized with Sac1p and Golgi-like liposomes enriched in PI and decorated with the PI 4-kinase. We established that Osh4p can create a sterol gradient between these membranes as soon as PI(4)P is synthesized following ATP addition, and determined the quantity of PI(4)P and ATP needed for this process. Third, we built a mathematical model which was validated by our *in vitro* data, and then was used to predict whether PI(4)P metabolism can pilot substantial ORP-mediated lipid transfer in cells. Finally, we showed that Sec14p, by transferring PI from ER- to Golgi-like membranes can support the synthesis of PI(4)P and the creation of a sterol gradient by Osh4p. These results improve our understanding on how ORPs distribute lipids under the control of PI(4)P metabolism in cells.

## Results

### PI(4)P synthesis on PI-rich liposomes by a membrane-bound PI 4-kinase

To reconstitute an ER/Golgi interface, a first step was to anchor Sac1 and a PI 4-kinase to distinct membranes. We already obtained a Sac1p[1–522]His_6_ construct that can be attached to liposomes, doped with 1,2-dioleoyl-*sn*-glycero-3-[(N-(5-amino-1-carboxypentyl)iminodiacetic acid)succinyl] (nickel salt) (DOGS-NTA-Ni^2+^), via an N-terminal His-tag that substitutes for the transmembrane segment of this protein (21). So we focused on the design of a PI 4-kinase that could be attached to liposomes using a distinct strategy. To this aim, we reengineered an active fragment of the PI-4 kinase IIα (PI4KIIα^SSPSS^ΔC, segment 78-453) in which the four cysteine residues of the ^174^CCPCC^178^ motif, which are normally palmitoylated to anchor the protein to membrane, have been replaced by serine residues to facilitate its purification (51). We re-introduced one cysteine in this motif (C178) and replaced three endogenous cysteine residues; exposed at the protein surface, by serine residues. Our idea was to covalently attach this kinase via this unique C178 residue to the surface of liposomes containing the thiol-reactive 1,2-dioleoyl-*sn*-glycero-3-phosphoethanolamine-N-[4-(p-maleimidophenyl)butyramide] (MPB-PE) lipid with an orientation allowing the recognition of the PI head group and its phosphorylation **(Fig.1A)**. This construct (thereafter called PI4K) was purified in the presence of DTT (**Fig. S1A**), then applied to a desalting column to remove DTT and incubated (at 4 µM) for 1 hour with liposomes (800 µM lipids) only composed of 1,2-dioleoyl-sn-glycero-3-phosphocholine (DOPC) or additionally containing 3 mol% MPB-PE, 10 mol% PI or both lipids. By flotation assays **(Fig.1B)**, we found that PI4K was weakly associated with pure DOPC liposomes (5 % of total protein) but seven-time more with MPB-PE-containing liposomes, which suggested that the protein was covalently attached to these latter. PI4K was slightly bound to liposomes doped with 10 mol% PI (10 % of total protein), possibly through a specific recognition of PI headgroup and/or non-specific electrostatic interaction. Finally, the highest degree of association (60 %) was measured with liposomes containing both PI and MPB-PE. These data suggested that PI4K can be efficiently anchored to liposomes enriched with PI.

**Figure 1.**
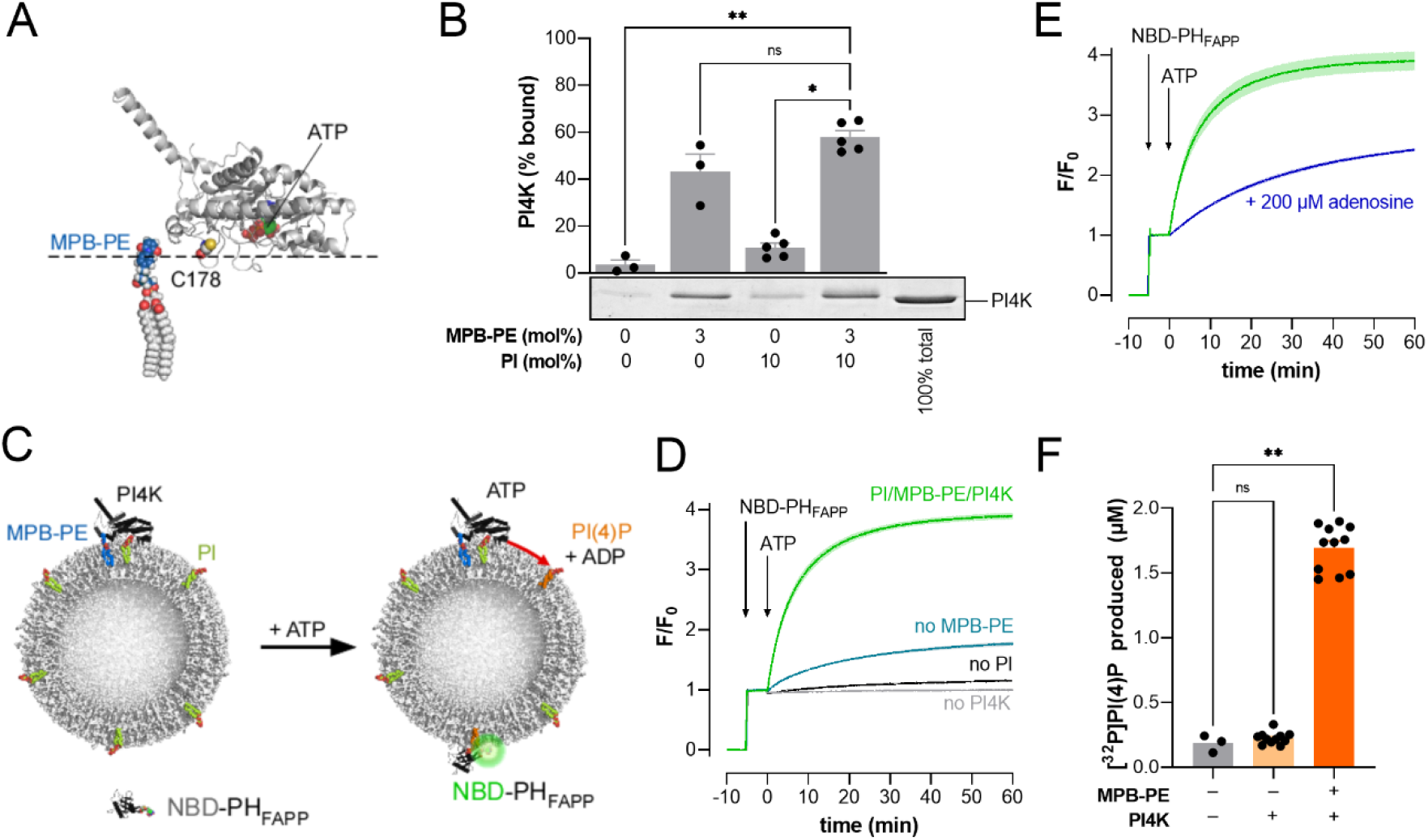
PI(4)P synthesis on liposome surface by membrane-associated PI 4-kinase. (A) Tridimensional structure of PI4KIIα^SSPSS^ΔC protein (PDB ID: 4HNE (51)). The solvent-exposed serine S178 in the SSPSS sequence was replaced by a cysteine residue (in sphere mode, with carbon in grey, nitrogen in blue, oxygen in red and sulfur in yellow). The molecule of ATP bound to the PI 4-kinase in the crystal is shown (in sphere mode, with carbon in green, nitrogen in blue, oxygen in red and phosphorus in orange). The orientation of the PI 4-kinase relative to the plane of the membrane (dashed line) has been suggested by molecular dynamics simulations (51). A molecule of MPB-PE is represented in sphere mode with carbon in grey (glycerol and acyl chain) or dark blue (headgroup), oxygen in red, nitrogen in blue, phosphate in orange and hydrogen in white. The figure was prepared using PyMOL (http://pymol.org/). (B) Flotation assays. PI4K (4 µM) was incubated in HK buffer at 25°C for 1 hour under agitation with liposomes (800 µM total lipids) only made of DOPC or containing 10 mol% liver PI, 3 mol% MPB-PE or both lipids. The functionalization reaction was stopped by adding 1 mM DTT. After centrifugation, the liposomes were recovered at the top of sucrose cushions and analyzed by SDS-PAGE. The amount of membrane-bound PI4K was determined using the content of lane 5 (100 % total) as a reference based on the SYPRO Orange signal. Data are represented as mean ± s.e.m (n = 3-5) with single data points. Kruskal–Wallis with Dunn’s multiple comparaison test (\**p* < 0.05, \*\**p* < 0.01). (C) Principle of PI(4)P detection in membrane decorated with PI4K using NBD-PH_FAPP_. ATP is added to PI-containing liposomes to which PI4K is covalently attached by MPB-PE lipid. NBD-PH_FAPP_ binds to PI(4)P that is synthesized in the outer leaflet of liposomes by the kinase, which results in an increase in its fluorescence signal. (D) Liposomes (800 µM lipids) composed of DOPC/liver PI/MPB-PE (87/10/3) were pre-incubated for 1 hour with 4 µM of DTT-free PI4K (PI4K/L molar ratio = 1/100 considering only accessible lipids in the outer leaflet). The mixture was then diluted 4-fold in a final volume of 600 µL HK buffer containing 7 mM MgCl_2_ (HKM7 buffer). After 5 minutes, once the sample reached the thermal equilibration at 30°C, NBD-PH_FAPP_ (250 nM) and ATP (100 µM) were sequentially injected. Control experiments were done in the absence of PI4K or with liposomes devoid of MPB-PE or PI. The signal was measured at λ = 525 nm (excitation at λ = 495 nm). The fluorescence is normalized to the F_0_ fluorescence recorded before the last injection. Each trace is the mean ± s.e.m. of several kinetics recorded in independent experiments (green trace, PI-containing liposomes decorated with PI4K, n = 4; blue trace, liposomes without MPB-PE, n = 3; dark trace, liposomes without PI, n = 3; gray trace, without PI4K, n = 4). (E) NBD-PH_FAPP_ and ATP were sequentially added to liposomes (200 µM lipids) composed of DOPC/liver PI/MPB-PE (87/10/3) and functionalized with PI4K (PI4K/L = 1/100). Measurement were done at 30°C in the absence or presence of 200 µM adenosine. Mean ± s.e.m. (green trace, without adenosine, n = 3; blue trace, with adenosine, n = 3). (F) Kinase assay. [γ-^32^P]ATP (100 µM) was added to DOPC liposomes (200 µM) containing 10 mol% PI, doped or not with 3 mol% MPB-PE, and pre-incubated for 1 hour with PI4K at PI4K/L= 1/200. Control experiments were done with liposomes containing 10 mol% PI in the absence of kinase. The reaction was let to proceed for 1 hour at 30°C under constant agitation. The lipids were extracted and dried. The radioactivity initially present in the reaction mix and associated to lipids was counted to determine the amount of synthesized PI(4)P. Dark orange bars, with MPB-PE-containing liposomes; light-orange bars, with liposomes devoid of MPB-PE, grey bars, without PI4K. Data are represented as mean ± s.e.m. (error bars; n *=* 3-11). Kruskal–Wallis with Dunn’s multiple comparison test; \*\**p* < 0.01, ns : *p* ≥ 0.05.

We then tested whether the kinase, attached to PI-rich liposomes, could synthesize PI(4)P in the presence of ATP **(Fig.1C)**. To this end, we followed in real-time the formation of PI(4)P in the outer leaflet of these liposomes using NBD-PH_FAPP_, a sensor whose fluorescence increases when it docks onto membrane by interacting with PI(4)P (21, 24). Liposomes composed of DOPC/PI/MPB-PE (87/10/3) were incubated with PI4K for 1 hour to be functionalized with the kinase and then diluted in a cuvette thermostated at 30°C. The final concentrations were 1 µM PI4K and 200 µM lipids (PI4K/lipid molar ratio = 1/100 considering only accessible lipids). NBD-PH_FAPP_ was then added, resulting in a signal jump at 525 nm (excitation at 495 nm). Five minutes after, ATP was injected (100 µM final concentration), which elicited a rapid increase in fluorescence, indicative of the progressive recruitment of NBD-PH_FAPP_ on liposomes **(Fig.1D)**. A plateau was reached with 1 hour. If the kinase was incubated with liposomes containing PI but not MPB-PE, the increase in NBD signal was much lower whereas with liposomes devoid of PI or kinase, a very slight and no augmentation were seen, respectively. Jointly these data suggested that PI4K, when it is covalently attached to PI-rich liposomes, generate PI(4)P.

We performed further experiments with PI-rich liposomes functionalized with PI4K in the absence or presence of 200 µM adenosine, an inhibitor of class II PI 4-kinases (52). As shown in **Fig. 1E**, the increase in signal was significantly slower in the presence of adenosine, confirming that our measurements did reflect a kinase-dependent PI(4)P synthesis process. Finally, we measured the production of PI(4)P by an independent way, using a radioactivity-based assay adapted from (53). PI-containing liposomes, doped or not with 3 mol% MPB-PE, were incubated with PI4K for 1 hour and then mixed with [γ-^32^P]ATP for an extra hour at 30°C. Control experiments with PI-rich liposomes in the absence of kinase were conducted. After liquid extraction, the amount of radioactivity incorporated in organic solvent-extractable material, resulting from the phosphorylation of PI, was counted. A significant level of radioactivity, corresponding to the production of ∼ 2 µM PI(4)P was measured, if PI4K was attached to liposomes containing MPB-PE **(Fig. 1F)**. In comparison, a negligible amount of radioactivity was measured both in experiments with liposomes devoid of MPB-PE and mixed with PI4K, or in the absence of PI4K. Jointly these data confirmed that the PI4K construct, attached to PI-rich liposomes, generated PI(4)P.

### Quantification of PI(4)P generated by the PI 4-kinase on liposome surface

Next we estimated how much PI(4)P was synthesized by PI4K through the analysis of the fluorescence signal. The increase in this signal cannot follow linearly the amount of PI(4)P that is synthesized. Indeed, it depends on a 1:1 binding process between NBD-PH_FAPP_ and PI(4)P that is governed by the affinity of the sensor for its ligand. Thus, to interpret our kinetics, we measured in separate experiments, the fluorescence of NBD-PH_FAPP_ in the presence of different liposomes each with a composition corresponding to a given degree of PI-to-PI(4)P conversion. In these liposomes, the fraction of PI(4)P ranged from 0 to 7.5 mol% of lipids at the expense of PI (so that PI and PI(4)P always represented a total of 10 mol% as in the kinase assays). The fluorescence was recorded with measurement settings identical to those used to follow PI(4)P synthesis. By plotting the intensity of NBD signal as a function of PI(4)P concentration, we obtained a binding curve that could be easily fitted assuming a 1:1 protein-ligand interaction model (**Fig.2A**). Then, using the obtained parameters (F_0_, F_max_, K_D_) and considering that a PI-to-PI(4)P conversion occurred following a pseudo first-order reaction, we fitted the data points of fluorescence traces measured in the presence of liposomes (200 µM lipids, 10 µM PI), decorated with 0.25, 5 or 1 µM PI4K (**Fig. 2B, Fig. S2A**) to estimate the synthesis of PI(4)P over time as a function of PI4K density on liposome surface (see Methods). The amount of PI(4)P synthesized after 1 hour was found to be proportional to this factor and to reach a maximum of 2.6 ± 0.4 µM PI(4)P (n = 4, ± s.e.m.) when liposomes were functionalized with a PI4K/lipid ratio of 1/100 (**Fig. 2D & Fig.S2A**). ORPs can extract PI(4)P from membrane with a 1:1 stoichiometry (21, 24, 54) and therefore be used to estimate how much of this lipid is present in a membrane. Thus, at the end of experiments done at PI4K/L = 1/100, we added incremental amounts of Osh4p and observed that the NBD signal returned to its initial level once 2.5-3 µM of Osh4p were present **(Fig. 2C)**. This suggested that a corresponding amount of PI(4)P was in liposome membrane and extracted by Osh4p, confirming our evaluation of PI(4)P production from kinetics curves. Next, we examined whether the PI4K activity could be influenced by the fluidity of the membrane, given that PI4KIIα might be sensitive to this factor, (55), by measuring its activity on membranes composed of POPC, instead of DOPC, and doped with PI and MPB-PE. We estimated that the production of PI(4)P was not higher **(Fig. 2D, Fig.S2B&C)**. On the contrary, we observed a faster production of PI(4)P when liver PI was replaced by soy PI, with liposomes functionalized with the same amount of PI4K **(Fig. 2D, Fig.S2D&E)**. This is possibly because these PI species differ in terms of their acyl chains (18:0/18:2 *vs* 18:0/20:4). This observation allowed us to use less PI4K to produce PI(4)P in some of the next assays.

**Figure 2.**
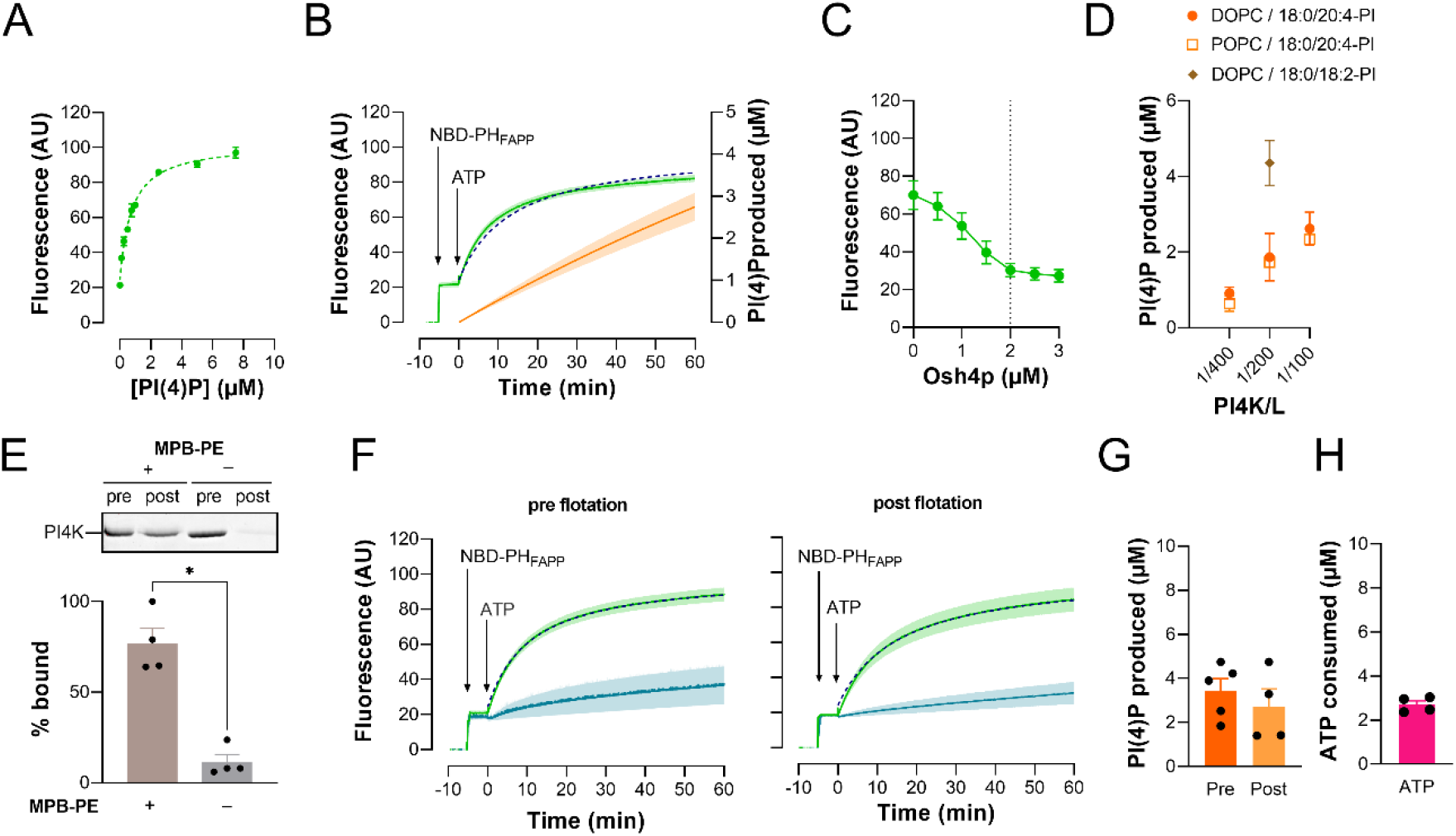
Quantification of PI(4)P production by membrane-anchored PI4K. (A) Fluorescence of NBD-PH_FAPP_ (250 nM) incubated with liposomes (200 µM lipids) made of DOPC and containing increasing amount of PI(4)P at the expense of PI (with PI + PI(4)P accounting for 10 mol% of total lipids), in HKM7 buffer at 30°C. The signal was measured with settings that were identical to those used for following PI(4)P synthesis. The intensity F is given as a function of the volume concentration of PI(4)P. This results in a binding curve that can be fitted considering F = F_0_ + F_B_ × [PI(4)P]/([PI(4)P] + K_D_) and a 1:1 stoichiometry. (B) Determination of PI(4)P production. Liposomes (800 µM lipids), composed of DOPC/liver PI/MPB-PE (87/10/3) and functionalized with PI4K (PI4K/L= 1/100), were diluted 4-fold in 600 µL of HKM7 buffer. Then, NBD-PH_FAPP_ (250 nM) and ATP (100 µM) were added. The fluorescence signal was measured at 525 nm (λ_ex_ = 495 nm, green trace) and corrected for photobleaching. The data points were fitted (dotted curve) to determine the k_syn_ rate of PI(4)P synthesis (considering a pseudo first-order reaction, see Methods). The amount of PI(4)P produced over time by PI4K was then calculated (orange curve). Mean ± s.e.m. (n = 3-4). (C) At the end of kinetics shown in (B), incremental amounts of Osh4p were added to liposomes in which PI(4)P was synthesized. The decrease in fluorescence indicated that Osh4p extracted PI(4)P. A minimal signal was reached once 2.5-3 µM Osh4p in total was added (dashed lines), suggesting that a similar amount of PI(4)P was present in the outer leaflet of liposomes. (D) Amount of PI(4)P synthesized after 1 hour in DOPC or POPC membranes enriched with 10 mol% liver PI (mostly 18:0/20:4 species) or 10 mol% soy PI (mostly 18:0/18:2 species), and functionalized with variable quantity of PI4K. Data are represented as mean ± s.e.m. (n = 3-5). (E) Purification of liposomes functionalized with PI4K. PI-containing liposomes (800 µM lipids), containing or not 3 mol% MPB-PE were mixed for 1 hour with 4 µM PI4K and then recovered at the top of sucrose cushions after centrifugation. SDS-PAGE analysis indicated that most kinase was attached to MPB-PE-containing liposomes (compare “pre” and “post” lanes, + MPB-PE). In contrast, almost no kinase was associated with MPB-PE-free liposomes (– MPB-PE) after the purification step. Each lane contains the same amount of liposome. Data are represented as mean ± s.e.m. (n = 4). Unpaired Mann–Whitney U test; \**p* < 0.05. (F) A 150-µL volume of a suspension of PI-containing liposomes, doped or not with MPB-PE, and mixed with PI4K, without further purification, was diluted 4-fold in 600 µL of HKM7 buffer. Alternatively, a 100-µL volume of the same liposome preparation, once purified by flotation as show in (E), was diluted 6-fold in 600 µL of HKM7 buffer. In each case, the final lipid concentration was 200 µM. NBD-PH_FAPP_ and ATP were added and PI(4)P production was followed by measuring NBD fluorescence. Mean ± s.e.m. (green trace, with liposomes doped with MPB-PE, n = 4-5; gray trace, liposome without MPB-PE, n = 4). (G) Amount of PI(4)P produced in PI-containing membranes, doped with MPB-PE and functionalized with PI4K within 1 hour following ATP addition, estimated by fitting the kinetics traces shown in (F). Data are represented as mean ± s.e.m. (orange bars, non-purified liposomes, n = 5; red bars, liposomes isolated by flotation, n = 4). (H) Amount of ATP specifically hydrolysed in 1 hour by PI4K at the surface of MPB-PE-containing liposomes used for assays shown in (F). Data are represented as mean ± s.e.m. (error bars; n = 4).

We further characterized the activity of PI4K by quantifying how much ATP was hydrolyzed by the kinase during PI(4)P synthesis. Preliminary assays showed that hydrolysis occurred in samples in which the PI4K was mixed with liposomes devoid of PI, possibly due to minute amounts of *E.coli* chaperones in protein preparation. To circumvent this problem, we devised a protocol in which liposomes composed of DOPC/soy PI/MPB-PE (87/10/3) were mixed with PI4K (PI4K/L = 1/200) for 1 hour, and then collected at the top of a sucrose gradient after ultracentrifugation to separate them from potential soluble contaminants. Similarly, we purified control liposomes devoid of MPB-PE but mixed with the same amount of PI4K. We estimated by SDS-PAGE analysis that 76.9 ± 8.4 % of PI4K were attached to MPB-PE-containing liposomes (**Fig. 2E**, n = 4, s.e.m.). Same results were obtained by measuring the intrinsic fluorescence of PI4K associated with liposomes before and after the purification step **(Fig. S2F)**. Using our NBD-PH_FAPP_-based assays, we estimated that 2.7 ± 0.8 µM PI(4)P were produced in the outer leaflet of purified liposomes (200 µM lipids) after a 1-hour incubation with ATP **(Fig. 2F,G)**. This was slightly less than the amount of PI(4)P measured for the same preparation of liposomes before centrifugation (3.5 ± 0.5, n = 4). In contrast, only 11.4 ± 4.1 % PI4K were bound to purified liposomes devoid of MPB-PE and almost no PI(4)P synthesis was measured **(Fig. 2F)**. In parallel, using a standard luminescence-based assay, we determined under the same conditions the amount of ATP hydrolyzed in 1 hour in each sample, i.e., with purified liposomes covered or not by kinase **(Fig. S2G)**. After data correction, we estimated that 2.7 ± 0.2 ATP molecules were specifically hydrolyzed by the kinase during this period of time **(Fig. 2H)**. This indicates that ∼1 molecule of ATP was used by PI4K to produce 1 molecule of PI(4)P, suggesting that PI(4)P synthesis occurred with no ATP-consuming futile cycle.

### PI(4)P synthesis and hydrolysis drive the generation of a sterol gradient between membranes by Osh4p

Once the PI4K activity was characterized, we set-up a full system to measure the sterol/PI(4)P exchange activity of Osh4p between two membranes, mimicking the ER/Golgi interface, under the control of a self-generated PI(4)P gradient. As a first step we assessed whether Osh4p could transfer PI(4)P synthesized by PI4K associated with a first population of liposome (Golgi mimetic, termed L_G_) to a second one (ER mimetic, L_E_), with a FRET-based assay using NBD-PH_FAPP_. L_E_ liposomes, made of DOPC (200 µM lipids) were mixed with an equal amount of L_G_ liposome, decorated with PI4K and doped with 10 mol% liver PI and 2 mol% 1,2-dipalmitoyl-*sn*-glycero-3-phosphoethanolamine-N-(lissamine rhodamine B sulfonyl) (Rhod-PE). We reasoned that the production of PI(4)P would trigger the association of NBD-PH_FAPP_ to L_G_ liposomes but the fluorescence of the sensor would barely or weakly increase, due to a FRET process with Rhod-PE present in these liposomes. However, we expected that, if newly-made PI(4)P was subsequently transported by Osh4p to L_E_ liposomes, this would provoke the translocation of NBD-PH_FAPP_ onto membranes devoid of Rhod-PE, and consequently an increase in fluorescence **(Fig. 3A).** Indeed, when ATP only was added to liposomes, the signal of the PI(4)P sensor did not change. On the contrary, when ATP and Osh4p (200 nM) were added together, the NBD signal progressively increased, suggesting that PI(4)P, once it was synthesized in L_G_ liposomes was transferred to L_E_ liposomes **(Fig. 3A)**. We repeated this experiment with L_E_ liposomes containing 2.5 mol% of dehydroergosterol, (DHE) which is a perfect ergosterol mimetic that is recognized by Osh4p (19, 21). The NBD signal increased more rapidly, which suggested that PI(4)P was delivered much faster to L_E_ liposomes. This was in line with the fact that the presence of DHE enhances the delivery of PI(4)P in membrane as Osh4p can exchange PI(4)P for this sterol (21). Jointly these data indicated that the PI(4)P pool generated by PI4K in L_G_ liposomes could be transferred by Osh4p to L_E_ liposomes and possibly exchanged with sterol contained in these liposomes.

**Figure 3.**
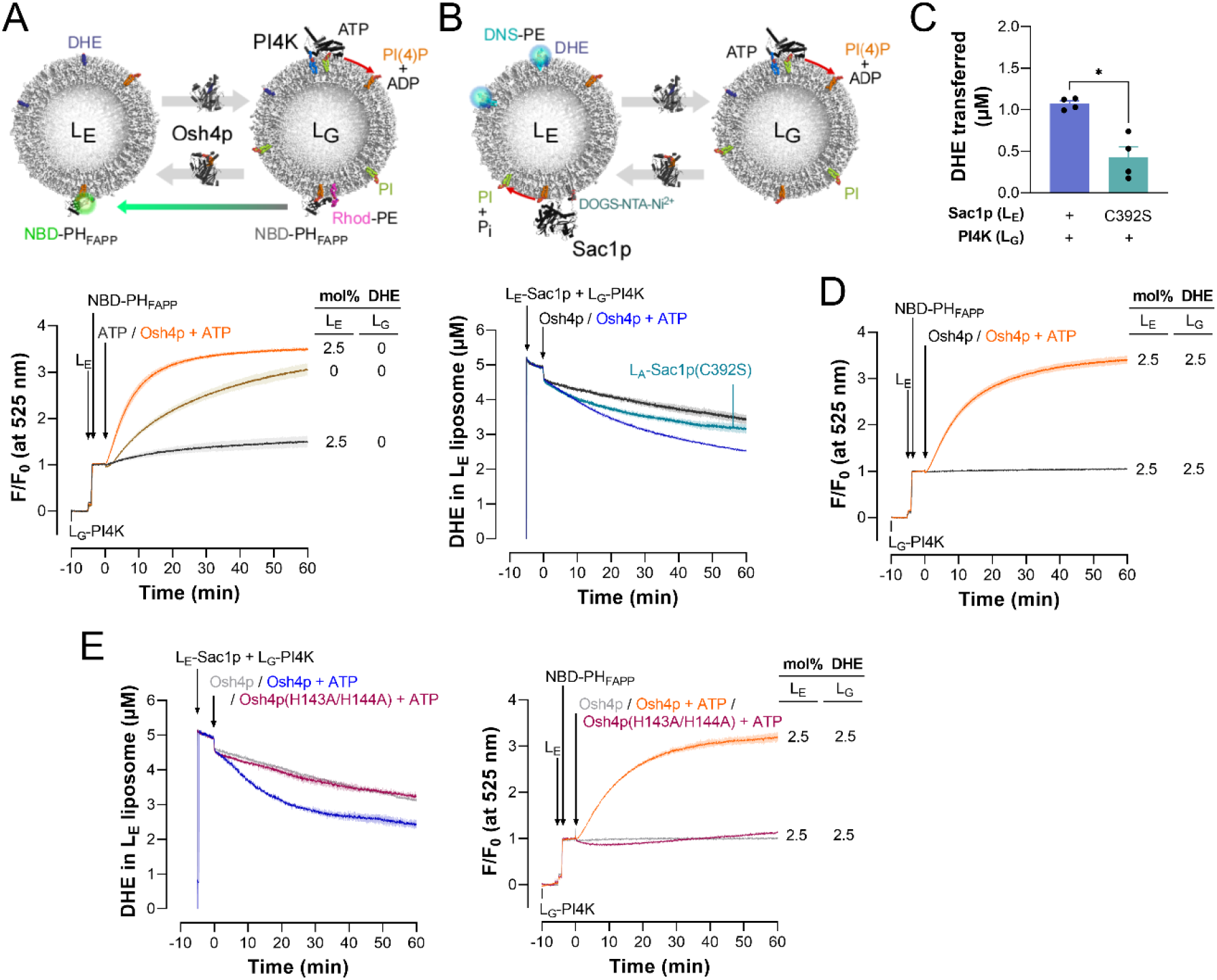
PI(4)P synthesis and hydrolysis drive sterol transfer by Osh4p between membranes. (A) Osh4p-mediated PI(4)P transfer coupled to PI(4)P synthesis. L_E_ liposomes (200 µM lipids) only made of DOPC, or additionally containing 2.5 mol% DHE, were added to an equal amount of L_G_ liposomes composed of DOPC/liver PI/MPB-PE/Rhod-PE (85/10/3/2) and functionalized with PI4K (at PI4K/lipid = 1/100) in 600 µL of HKM7 buffer at 30°C. One minutes after, NBD-PH_FAPP_ (250 nM) was added. After a 5-min incubation, ATP only (100 µM) or together with Osh4p (200 nM) was injected. Fluorescence is measured at 525 nm (λ_ex_ = 460 nm). Mean ± s.e.m. (grey trace, ATP without Osh4p, n = 3; brown and orange trace, ATP pre-mixed with Osh4p, n = 3-4). (B) Osh4p-mediated DHE transfer driven by a PI(4)P gradient generated by PI4K and Sac1p. L_E_ liposomes (200 µM lipids), composed of DOPC/DHE/DNS-PE/DOGS-NTA-Ni^2+^ (93/2.5/2.5/2), and covered with Sac1p[1-522]His_6_ or its C392S inactive form (100 nM), were added to an equal amount of L_G_ liposomes composed of DOPC/liver PI/MPB-PE/DHE (84.5/10/3/2.5) and functionalized with PI4K (PI4K/L = 1/100) in HKM7 buffer. After a 5-min incubation at 30°C, 200 nM Osh4p, alone or together with 100 µM ATP, were injected. Fluorescence was recorded at 525 nm (λ_ex_ = 310 nm) and normalized in terms of DHE present in L_E_ liposomes. Mean ± s.e.m. (grey trace, Osh4p without ATP, L_E_ liposomes decorated with Sac1p[1-522]His_6_, n = 4; blue trace, Osh4p pre-mixed with ATP, L_E_ liposomes decorated with Sac1p[1-522]His_6_, n=4 ; blue-green trace, Osh4p pre-mixed with ATP, L_E_ liposomes decorated with Sac1p[1-522]His_6_(C392S), n = 4). (C) Amount of DHE specifically transferred by Osh4p from L_E_ liposomes, decorated with active Sac1p (blue bar) or its inactive C392S form (green bar), to L_G_ liposomes covered with PI4K within 1 hour in the presence of ATP. Data are represented as mean ± s.e.m. (error bars; n = 4). Unpaired Mann– Whitney U test; \**p* < 0.05. (D) PI(4)P-transport coupled to PI(4)P synthesis between L_E_ and L_G_ liposomes both containing 2.5 mol% DHE. L_G_ liposomes (200 µM lipids) composed of DOPC/liver PI/MPB-PE/DHE (84.5/10/3/2.5) and functionalized with PI4K (PI4K/L = 1/100) were mixed with an equal amount of L_E_ liposomes, composed of DOPC/DHE (97.5/2.5) in HKM7 buffer. NBD-PH_FAPP_ was added (250 nM) followed by the injection of 200 nM Osh4p, alone or with 100 µM ATP. Fluorescence was measured at 525 nm on excitation at 495 nm. Mean ± s.e.m. (grey trace, Osh4p without ATP, n = 3; orange trace, Osh4p pre-mixed with ATP, n = 4). (E) PI(4)P synthesis cannot drive the transfer of DHE by Osh4p(H143A/H144) mutant. Left panel: experiments were done as in (B) with L_E_ liposome decorated with Sac1p and L_G_ liposome containing 10 mol% soy PI. A final concentration of Osh4p or Osh4p(HH/AA) alone (200 nM), or together with 100 µM ATP was injected. Right panel: experiments were done as in (D) with 200 nM Osh4p or Osh4p(H143A/H144A), and L_G_ liposomes doped with 10 mol% soy PI.

Next, we modified our system to measure whether Osh4p could transfer sterol from L_E_ to L_G_ membranes in the presence of a PI(4)P gradient generated between these membranes by PI4K and Sac1p (**Fig. 3B**). We prepared L_E_ liposomes containing 2.5 mol% DHE and DNS-PE (1,2-dioleoyl-*sn*-glycero-3-phosphoethanolamine-N-(5-dimethylamino-1-naphthalenesulfonyl)) but also 2 mol% DOGS-NTA-Ni^2+^. They were functionalized with Sac1p[1–522]His_6_ (100 nM) or with a catalytically-dead version of this construct (C392S). In parallel, we prepared L_G_ liposomes composed of DOPC/PI/DHE (87.5/10/2.5) covered by PI4K. Prior to each measurement of sterol transfer, the ability of PI4K to generate PI(4)P in DHE-containing membranes was verified (**Fig. S3A)**. Moreover, by flotation assays, we established that there was no exchange of Sac1p and PI4K between L_E_ and L_G_ liposomes, when these liposomes were mixed together **(Fig. S3B, S3C)**; this meant that the spatial separation of the two enzymes, necessary for the creation of a PI(4)P gradient between membranes, could be maintained over time. To quantify the transfer of DHE from L_E_ to L_G_ liposomes by Osh4p, equivalent amounts of L_E_ and L_G_ liposome (200 µM lipids each) were mixed, and the FRET signal between DHE and DNS-PE contained in L_E_ membrane was measured. Adding Osh4p alone (200 nM) to liposomes only provoked a small drop in signal followed by a slow decrease that was identical to that measured before injecting the protein, likely due to photobleaching (**Fig. 3B**, gray trace). This suggested that Osh4p only extracted a stoichiometric amount of DHE from L_E_ liposomes with no transfer to L_G_ liposomes. In contrast, when ATP was added to trigger PI(4)P synthesis, along with Osh4p, a faster decrease in signal was observed, indicating that Osh4p delivered sterol to L_G_ liposomes **(Fig. 3B**, blue trace). Importantly, given that DHE was initially equilibrated between membranes, this indicated that Osh4p was able to create a sterol gradient once PI(4)P was synthesized. Interestingly, when L_E_ liposomes were decorated with active Sac1p, the amount of transferred DHE after 1 hour was higher than in the presence of its catalytically-dead form (1.1 ± 0.03 µM vs 0.42 ± 0.13 µM (n = 4), **Fig. 3B&C**). Parallel assays using NBD-PH_FAPP_ showed that Osh4p transferred PI(4)P, synthesized in L_G_ liposomes in the presence of ATP, to L_E_ liposomes (covered by inactive Sac1p so that PI(4)P was detectable), with both types of liposome initially containing 2.5 mol% DHE as in the DHE transfer assays **(Fig. 3D)**. To strengthen our data, we repeated the DHE and PI(4)P transfer experiments with an Osh4p(H1443A/H144A) mutant deficient in binding PI(4)P (19, 21). As expected, we found that this protein was unable to transfer sterol and PI(4)P between membranes when PI(4)P was synthesized **(Fig. 3E)**. Jointly these data indicate that Osh4p can carry sterol from the ER to the Golgi mimetic membranes, in which PI(4)P is synthesized, and transport this PI(4)P pool in the opposite direction. This means that the production of PI(4)P alone can drive the creation of a sterol gradient between the two membranes but that PI(4)P hydrolysis on the ER mimetic membrane markedly improves this process. These results demonstrate that, in the presence of ATP, distant kinase and phosphatase can generate a PI(4)P gradient between membranes and thereby, drive the creation of a second lipid gradient by ORP-mediated lipid exchange.

### Relationship between ATP-dependent PI(4)P synthesis and sterol transfer

Next, we quantified how much PI(4)P was synthesized to allow for the creation of a sterol gradient between membranes by Osh4p. L_G_ liposomes composed of DOPC/soy PI/DHE/MPB-PE (84.5/10/2.5/3) were functionalized with two different concentrations of PI4K (PI4K/L = 1/200 or 1/100). Then, with each liposome preparation, we carried out two parallel measurements. First, we determined the production of PI(4)P by the PI4K within 1 hour using the NBD-PH_FAPP_ sensor. Secondly, we mixed the L_G_ liposomes with L_E_ liposomes composed of DOPC/DHE/DNS-PE (95/2.5/2.5) covered by Sac1p, and quantified the transfer of DHE between these liposomes by Osh4p in the presence of ATP; control experiments were carried out with kinase-free L_G_ liposomes. The production of PI(4)P after one hour was higher if L_G_ liposomes were functionalized with a higher density of PI4K **(Fig. S3D)**. In parallel, we measured that more DHE was transferred from L_E_ to L_G_ liposomes if more kinase was used **(Fig. 4A)**. We established that the synthesis of 2.6 and 6.7 µM PI(4)P drove the transfer of 1 µM and 1.5 µM DHE, respectively, indicating that the transfer of one molecule of DHE during the creation of a sterol gradient required the synthesis of more than one PI(4)P molecule **(Fig. 4B, Fig.S3D)**. This analysis also indicated that, for the generation of a steeper sterol gradient, a higher production of PI(4)P was needed for the transfer of a similar DHE amount **(Fig. 4C)**. Overall, these data suggested that the creation of a sterol gradient did not simply result on a one-for-one exchange of sterol and PI(4)P in our system. To better analyze this, we tested whether a pre-established DHE gradient between L_E_ and L_G_ membranes could be dissipated through spontaneous DHE transfer in the time-scale of our assays. We mixed L_E_ liposomes containing 1.5 or 0.5 mol% DHE with L_G_ liposomes containing 3.5 or 4.5 mol% DHE, respectively, in the absence of protein and measured a substantial transfer of DHE from L_G_ to L_E_ liposomes in one hour (up to 1.6 µM DHE with the steepest gradient, **Fig. S3E**). We concluded that if the creation of a sterol gradient by Osh4p required a PI(4)P amount higher than expected, this was likely because spontaneous DHE transfer counteracted the creation of this gradient.

**Figure 4.**
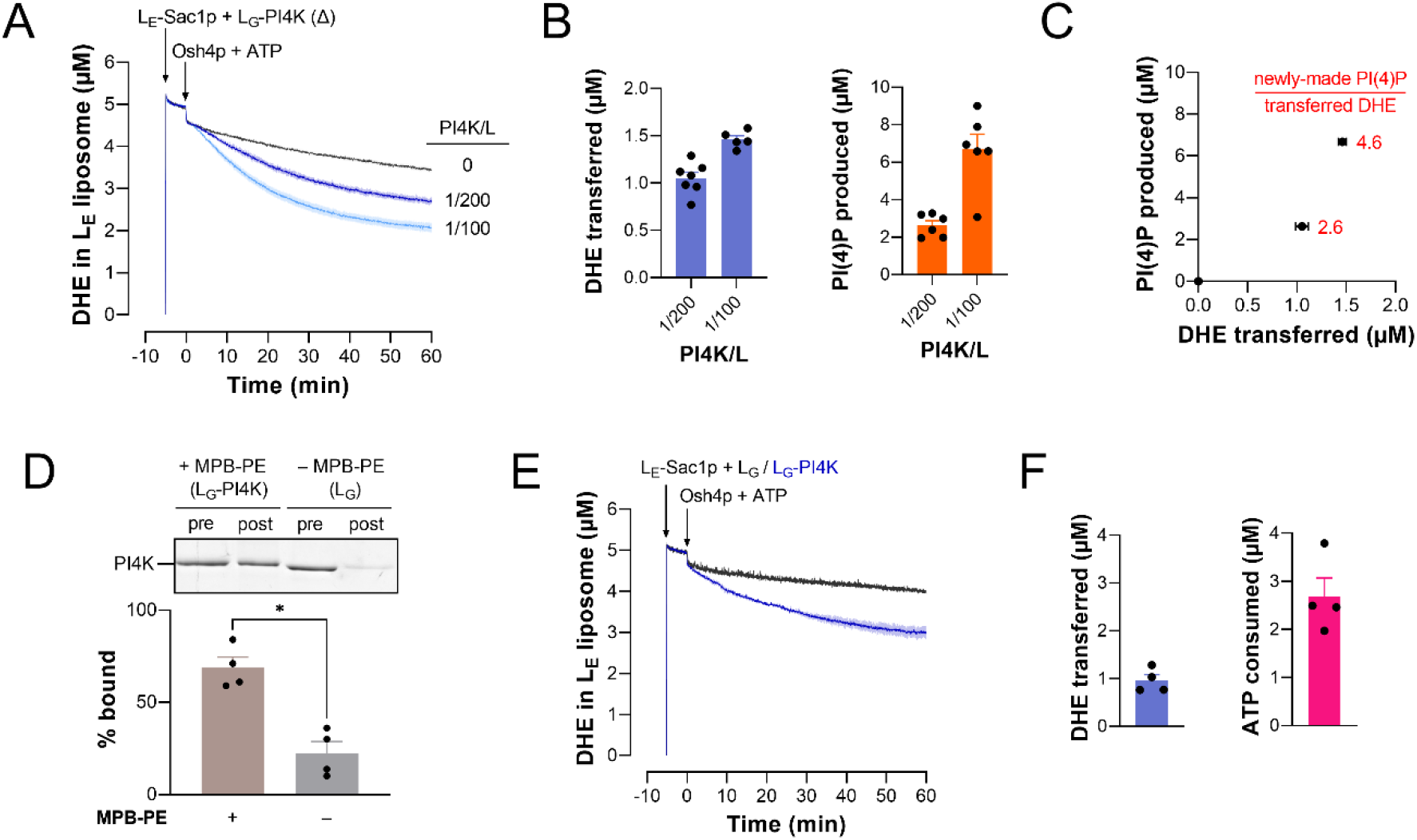
Relationship between PI(4)P synthesis and Osh4p-mediated sterol transfer. (A) Osh4p-mediated DHE transfer as a function of PI4K density on L_G_ liposomes. L_E_ liposomes (200 µM lipids) composed of DOPC/DHE/DNS-PE/DOGS-NTA-Ni^2+^ (93/2.5/2.5/2) covered with Sac1p[1-522]His_6_ (100 nM) were added to L_G_ liposomes (200 µM) composed of DOPC/soy PI/MPB-PE/DHE (84.5/10/3/2.5) and functionalized with different amounts of PI4K (PI4K/L= 0, 1/100 or 1/200). After five minutes, Osh4p (200 nM) and ATP (100 µM) were injected. Mean ± s.e.m. (n = 6-7). (B) Amount of DHE transferred by Osh4p from L_E_ to L_G_ liposomes and amount of PI(4)P produced in L_G_ liposomes in 1 hour in the presence of ATP, as a function of the quantity of PI4K used to functionalize L_G_ membranes. Data are represented as mean ± s.e.m. (error bars; n = 6) (C) Amount of Osh4p-mediated DHE transfer as a function of PI(4)P synthesized in L_G_ membrane by PI4K. Data are represented as mean ± s.e.m. (error bars) (D) Liposomes containing 2.5 mol% DHE, doped or not with 3 mol% MPB-PE, were mixed with PI4K (PI4K/L = 1/200) and purified by flotation. SDS-PAGE analysis indicates that most of the kinase was attached to MPB-PE-containing liposomes. Each lane contains the same amount of liposome. Data are represented as mean ± s.e.m. (error bars; n = 4). Unpaired Mann–Whitney U test; \**p* < 0.05. (E) DHE transfer assay using purified L_G_ liposomes decorated with PI4K. A 100-µL volume of the suspension of liposomes containing or not MPB-PE, mixed with PI4K, and purified by flotation as shown in (E), was diluted in a final volume of 570 µL of HK buffer supplemented with 7.4 mM MgCl_2_. Then, a 30-µL volume of L_E_ liposomes suspension, composed of DOPC/DHE/DNS-PE/DOGS-NTA-Ni^2+^ (93/2.5/2.5/2) and covered with Sac1p[1-522]His_6_ were added. The final lipid concentration of each liposome population is 200 µM. Five minutes after, Osh4p (200 nM) and ATP (100 µM) were injected simultaneously. Mean ± s.e.m. (grey trace, L_G_ liposomes devoid of MPB-PE with no attached PI4K, n = 4; blue trace, L_G_ liposomes doped with MPB-PE and decorated with PI4K, n = 4) (F) Amount of DHE transferred by Osh4p from L_E_ liposomes to L_G_ liposomes decorated by PI4K (derived from curves shown in (I)) and corresponding amount of ATP specifically hydrolyzed by the kinase within 1 hour. To determine ATP consumption, the purified L_G_ liposomes, naked or decorated with PI4K, used in experiments shown in panel I, were diluted in 100 µL of HKM7 buffer (200 µM lipids, final concentration) and incubated with 100 µM ATP for 1 hour at 30°C under agitation. The amount of hydrolyzed ATP determined in condition where PI4K was free was subtracted from that measured with the membrane-bound kinase. Data are represented as mean ± s.e.m. (error bars; n = 4).

To extend our analysis, two types of L_G_ liposome, with or without MPB-PE, both enriched with soy PI and DHE, were mixed with kinase (at PI4K/L = 1/200) and purified on sucrose gradient. SDS-PAGE analysis indicated that 68.8 ± 5.7 % of PI4K were bound to MPB-PE-containing liposomes (**Fig. 4D**, n = 4, s.e.m.). Then, comparative DHE transfer experiments along with assays to measure ATP hydrolysis were conducted using each type of L_G_ liposomes (**Fig. 4E, F**). We determined that 0.96 ± 0.13 µM of DHE were transported in 1 hour by Osh4p from L_E_ liposomes to L_G_ liposomes, if these latter were functionalized with kinase. Over the same period of time, ∼2.6 µM of ATP were hydrolyzed by the kinase at the surface of L_G_ liposomes, a value that matched well with the amount of PI(4)P that had to be synthesized for the transfer of 1 µM DHE in the same exact condition **(Fig. 4C, D).** We concluded that the amounts of hydrolyzed ATP and newly-made PI(4)P necessary for the transfer of a given amount of DHE by Osh4p were similar, corroborating our observations that PI4K uses one ATP to produce one PI(4)P molecule **(Fig. 2G,H).**

### Kinetic model of Osh4-mediated sterol/PI4P exchange powered by PI(4)P metabolism

Next, we developed a kinetic model to analyze our *in vitro* data and then estimate to what extent PI(4)P metabolism can drive the creation of sterol gradient in a cell. We considered a minimal model that describes, production and hydrolysis of PI(4)P, and transport of PI(4)P and sterol *in vitro*. We defined six unknown concentrations: PI(4)P in ER-like membranes (*p*_*E*_), in Golgi-like membranes (*p*_*G*_) and in complexes with Osh4p (*p*_*o*_), and similarly for sterol (*s*_*E*_, *s*_*G*_ and *s*_*o*_, **Fig. 5A)**. We wrote 6 non-linear kinetic equations with a specific form that was justified experimentally, as described in Sup. Materials and Methods (equation (7)). In our DHE-transfer assays, all Osh4p proteins were initially pre-loaded with sterol, hence *s*_*o*_ = *O* (concentration of Osh4p) and *p*_*o*_ = 0. We integrated numerically the full system using Π = 1.9 × 10^−3^μM. s^−1^ corresponding to a production of 6.7 µM at PI4K/L = 1/100 in one hour (**Fig. 4B**); *k*_*sac*_ = 0.0028 *s*^−1^ corresponding to the rate of PI(4)P hydrolysis by 100 nM Sac1p (based on (21)) and *r* = 0.015 ± 0.0009 μM^−1^. s^−1^, corresponding to the rate of lipid exchange between Osh4p and membranes (obtained from experiments with Osh4 preloaded with PI(4)P and mixed with a single population of liposome containing different amounts of DHE (**Fig. S4**)). We found that, as the production of PI(4)P starts at the surface of L_G_ liposomes, *p*_*G*_ builds up, hence at L_G_ liposomes, Osh4p starts to pick up PI(4)P and drop its sterol cargo. Therefore, in this system, the net effect is that the increase in PI(4)P concentration in L_G_ membranes results in the transport of sterol from L_E_ to L_G_ membranes (**Fig. 5B**). To better analyze this result, let us discuss more precisely the basis for the kinetic model. We considered the following reaction on both kinds of liposomes:

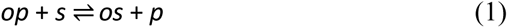

**Figure 5.**
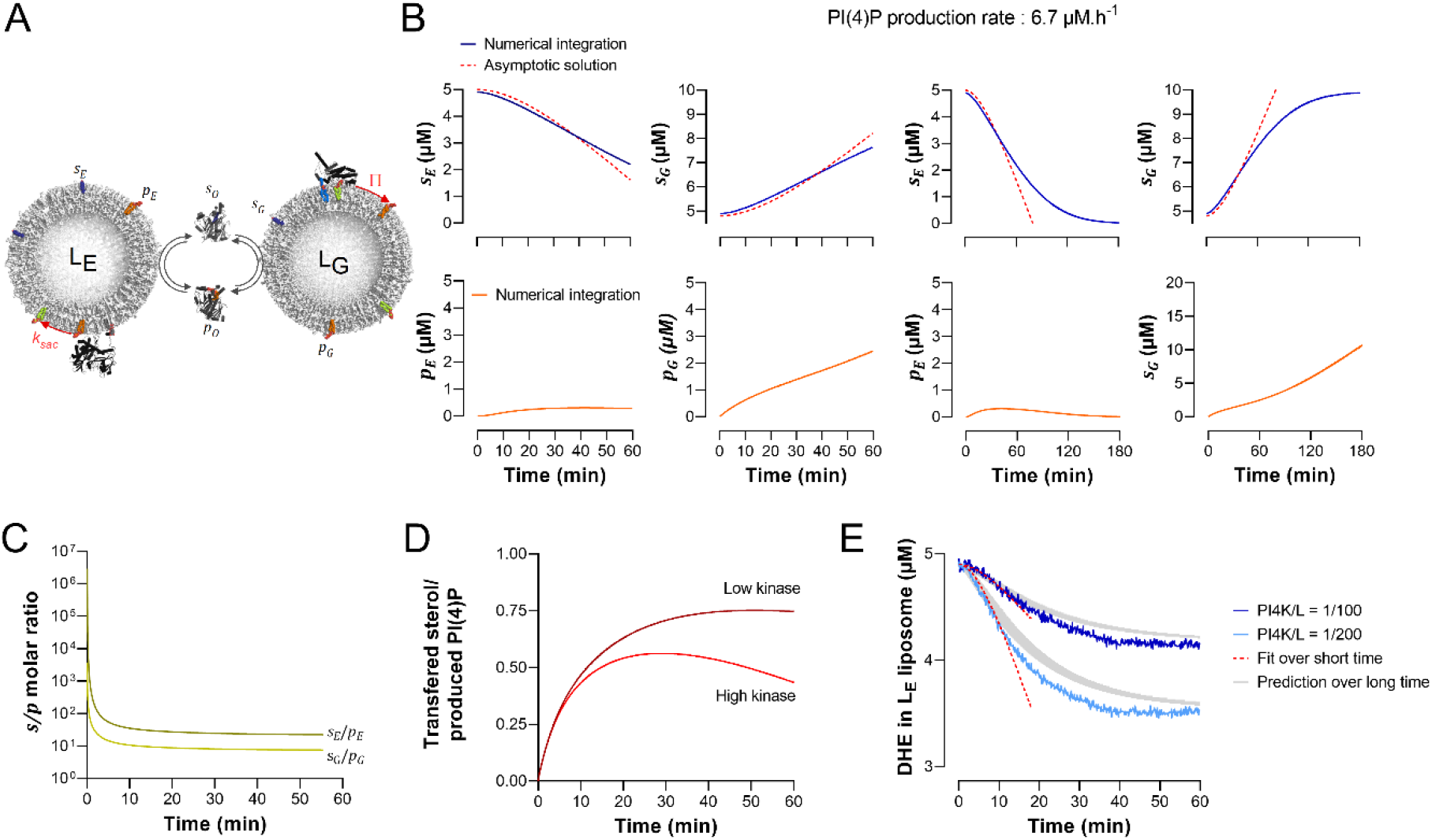
Kinetic model of ORP-mediated lipid exchange driven by PI(4)P metabolism. (A) Description of the minimal kinetic model (see details in the main text). (B) Evolution of sterol and PI(4)P concentration in the ER-like membrane (*s*_*E*_, *p*_*E*_) and Golgi-membrane (*s*_*G*_, *p*_*G*_) over a period of 1 h (left) or over a longer period of 3 h (right) according to the kinetic model. The concentration of Osh protein (*O*) is equal to 200 nM, the initial concentration of sterol *s*_*E*_ and *s*_*G*_ is equal to 5 µM and the PI(4)P synthesis rate Π = 1.9 × 10^−3^ µM. s^−1^, similar to the condition of DHE-based transfer assays (**Fig 4A** with PI4K/L = 1/100). The values were obtained by integrating numerically the full system (eq. (9) Materials and Methods) with numerical integration, (blue or orange) or by an asymptotic analysis of the full system (asymptotic solution, dashed red line). (C) Evolution of the *s*_*E*_/*p*_*E*_ and *s*_*G*_/*p*_*G*_ ratio over time in the system. (D) Evolution of the ratio of sterol transfer from ER-like to Golgi-like membrane to PI(4)P production in Golgi-like membrane with Π = 7 × 10^−4^ μM. s^−1^, or Π = 1.9 × 10^−3^ µM. s^−1^ . (corresponding to PI4K/L = 1/200 and 1/100, respectively). (E) Analysis of Osh4p-mediated DHE transfer curves shown in Fig. 4A (condition PI4K/L = 1/100 and 1/200) corrected for spontaneous signal decrease. The short term asymptotic were fitted to the first 100 data points (100 seconds) of the curve considering O = 200 nM and either Π = 7. 10^−4^ μM. s^−1^, or Π = 1.9. 10^−3^ µM. s^−1^ . A numerical integration of the full kinetic model, taking into account spontaneous sterol transfer between membrane provided quantitatively accurate predictions for the data at all times (grey shades: range of predictions for all admissible of *r* and *k*_*sac*_ value of the parameters from our fitting procedure).

Based on (19), we assumed that Osh4p has the same affinity for both lipids, thus forward and backward reaction rates are considered equal. Reaction (1) is at equilibrium in L_G_ membranes if *p*_*o*_ × *s*_*G*_ = *s*_*o*_ × *p*_*G*_. When *p*_*G*_ starts to build up, the reaction moves backward and thus *p*_*o*_ and *s*_*G*_ increase which corresponds to the notion that producing PI(4)P in L_G_ membranes drives transport of sterol to these membranes. In other words, as long as the system produces *p*_*G*_, *both p*_*G*_*and s*_*G*_*increase*. In L_E_ membrane, the opposite occurs. equation (1) is at equilibrium at L_E_ liposomes if *p*_*o*_ × *s*_*E*_ = *s*_*o*_ × *p*_*E*_, but on L_E_ liposomes Sac1p hydrolyzes PI(4)P, thus decreases *p*_*E*_ and reaction (1) moves forward, decreasing *s*_*E*_. As a result, on L_E_ liposomes, as long as Sac1p keeps depleting PI(4)P, *both p*_*E*_ *and s*_*E*_ *decrease.* The above argument shows that, in order for sterol to be transported from L_E_ liposomes to L_G_ liposomes, *p*_*o*_ × *s*_*G*_ < *s*_*o*_ × *p*_*G*_ and *p*_*o*_ × *s*_*E*_ > *s*_*o*_ × *p*_*E*_ hence:

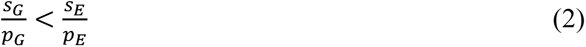

Thus, as long as the sterol-to-PI(4)P molar ratio is lower in L_G_ membranes than in L_E_ membranes, net sterol transfer occurs from L_E_ to L_G_ membranes. In other words, in this system, sterol is transported to L_G_ membranes if PI(4)P also increases in this membrane, even more than sterol. In our simulations, condition (2) was true as seen from **Fig. 5C**. A conclusion, perhaps counter intuitive, is that the maintenance of a low sterol-to-PI(4)P molar ratio in L_G_ membranes compared to L_E_ membranes is compatible with a transfer process that yields a higher *absolute* concentration of sterol in Golgi-like membranes than in ER-like membranes, *s*_*G*_ > *s*_*E*_.

Next, we observed that at longer times sterol transfer rate would increase and could eventually equal PI(4)P synthesis rate. The timescale to reach equilibrium between PI(4)P synthesis and sterol transport rates can be explained by an asymptotic solution of the full system, which is valid at short times (see **Fig. 5B** and **Fig. S5**). This short-term solution prescribes that, starting from *s*_*E*_(*t* = 0) = *s*_*G*_(*t* = 0) = *S*/2, sterol decrease in L_E_ membranes according to:

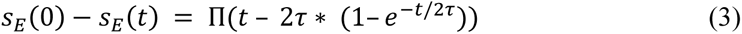

where

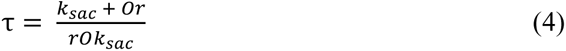

Equations (3) and (4) illuminate the role of PI(4)P production and hydrolysis for Osh4p-mediated sterol transfer: if Π increases, sterol transfer increases, according to equation (3), and if *k*_*sac*_ increases, *τ* decreases according to equation (4) which results in a faster sterol transfer (equation (3)). These equations also indicate that if the concentration of lipid exchanger and its exchange efficiency *r* increase, *τ* decreases (equation (4)) which results in a faster sterol transfer (equation (3)). To sum up, PI(4)P synthesis and hydrolysis as well as Osh4p-mediated lipid exchange amplitude speed up sterol transfer.

Numerical simulations showed that when *t* ≫ *τ*, the rate of sterol transfer increased, reaching almost the rate of PI(4)P synthesis Π (see **Fig. 5D**, with parameters chosen as described below). This means that a certain time is needed to observe that, for each molecule of PI(4)P produced in L_G_ membranes, one sterol gets transported to this membrane. Eventually, transport slows down and stops once all sterol is depleted from L_E_ membranes and accumulates entirely in L_G_ membranes, which was confirmed from our numerical simulations of the full system (**Fig. 5B**, over long time). Overall, the prediction from the kinetic model is that the synthesis of PI(4)P enables sterol transfer from L_E_ to L_G_ membranes, exactly as seen in our *in vitro* experiments. However, in our assays, a non-trivial equilibrium was reached *before* complete depletion of sterol in ER-membrane **(Fig. 4A)** which was inconsistent with the mathematical model. Likely, the reason for this is that spontaneous sterol transfer occurred between L_E_ and L_G_ liposomes (**Fig. S3E)**, counteracting the directional Osh4p-mediated transfer of sterol between these liposomes. Therefore, we included an additional term in the mathematical model that accounted for spontaneous sterol transfer (Sup. Materials and Methods, equation (16)).

Next, we directly compared theory and experimental results shown in **Fig. 4A**. To do so, we fitted the first 100 seconds of the DHE transfer kinetics, measured at PI4K/L = 1/100 and PI4K/L = 1/200 with the asymptotic solution, considering Π = 1.9 × 10^−3^ μM. s^−1^ and Π = 7.2 × 10^−4^ μM. s^−1^ respectively, corresponding to a production of 6.7 µM at PI4K/L = 1/100 in one hour and 2.6 µM at PI4K/L = 1/200 in one hour (**Fig. 4B**). This provided a fit for *τ* and thus estimates for *k*_*sac*_ = 0.015 ± 0.009 s^−1^ and *r* = 0.08 ± 0.05 μM^−1^. s^−1^, consistent with the *k*_*sac*_ and *r* values used previously. We then fitted the long-term equilibrium data to obtain the rate of spontaneous exchange, *C* = (4.7 ± 0.3) × 10^−4^ *s*^−1^ and *C* = (6.6 ± 0.2) × 10^−4^ s^−1^ at PI4K/L = 1/100 and PI4K/L = 1/200 respectively. Given these parameters, we solved numerically the full model to obtain quantitatively accurate predictions for the data at all times (grey shades in **Fig. 5E** representing the range of predictions for all admissible values of the parameters from our fitting procedure). Comparable results were obtained by analyzing the DHE transfer kinetics measured at PI4K/L = 1/200, indicating that our kinetic model explained well the *in vitro* data.

We next used this model, validated with *in vitro* experiments, to estimate whether the amount of PI(4)P generated in the Golgi membrane can maintain a proper sterol distribution at the ER/Golgi interface in a yeast or human cells. We assumed that cell membranes have relatively stable lipid composition, hence that lipid transport is at equilibrium. In that situation, for each PI(4)P molecule produced in the Golgi, membrane, one sterol molecule is transferred from the ER to the Golgi. Thus once the cell is at the desired equilibrium state, no PI(4)P is needed unless there is a depletion of sterol from the Golgi. In this case, PI(4)P production is needed to fuel further sterol transport which offsets this loss of sterol (see terms E, D in eq (17), Sup. Materials and Methods). Thus by tuning PI(4)P production, the cell can replenish the depleted sterol in the Golgi membrane, by increasing sterol transport from the ER membrane. However, PI(4)P production can only replenish sterol depletion up to a maximum amount that is dictated by the concentration of Osh4p (OSBP in human cell), O, the rate of Osh4p binding, r, and sterol concentration in the ER membrane (more details about the derivation of this bound can be found in Sup. Materials and Methods, in the discussion that leads to equation 22):

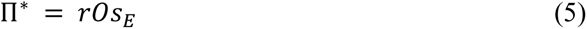

The actual sterol transport rate, Π, is smaller than Π^∗^ and could be estimated from knowledge of *S*, *O*, *s*_*E*_, *s*_*G*_, *p*_*E*_, *p*_*G*_, *k*_*sac*1_, *r* (see equation (23), Sup. Materials and Methods). We thus carefully collected information about the volume of a single yeast and HeLa cell, the total area of the ER and Golgi membrane in these cell types, sterol and PI(4)P levels in these membranes, and number of Osh4p/OSBP and Sac1p/SAC1ML copies *per* cell, from available studies (*see* Material and Methods). From these values, we estimated that in a yeast cell : *O* = 0.67 μM, *s*_*E*_ = 337 μM, *s*_*G*_ = 413 μM (**Table S1**). The, we used our estimates for the rate of lipid exchange *in vitro*, *r* = 0.08 μM^−1^. s^−1^, and for the rate of PI(4)P hydrolysis *k*_*sac*_ = 0.006 s^−1^ (see note **Table S1).** Plugging these values into equation (23) (Sup. Materials and Methods), we obtained a rate of ER-to-Golgi sterol transport Π = 0.25 μM. s^−1^ (**Table S1)**. Because at equilibrium the rate of PI(4)P synthesis at the Golgi equals the rate of PI(4)P hydrolysis of PI(4)P at the ER and the rate of ER-to-Golgi sterol transfer, all of these rates are expected to be on the order of 0.25 μM. s^−1^ i.e. 900 μM in one hour. This figure is compatible with the fact that in yeast devoid of Sac1p, PI(4)P level increases fivefold within an hour (37), which corresponds to about 300 µM, using our estimated Golgi PI(4)P concentration ≈ 62.8 μM, suggesting that Sac1p can hydrolyze about 240 μM PI(4)P *per* hour **(Table S1)**. Importantly, this estimation about PI(4)P hydrolysis and our predictions suggest that Osh4p coupled to PI(4)P metabolism can substantially renew the Golgi sterol pool (∼ 400 µM) in one hour.

For HeLa cells, we estimated that *O* = 0.364 μM, *s*_*G*_ = 404 μM; for the concentration of sterol in the ER membrane, we considered two extreme scenarios – either OSBP has only access to a pool of sterol present in ER patches engaged in contact sites with the Golgi or to sterol contained in the whole ER membrane – and obtained different estimates *s*_*E*_ = 67.5 or 844 μM. Consequently, we obtained two predictions for the rate of ER-to-Golgi sterol transport Π = 0.42 or 4 μM. s^−1^ which are, assuming equilibrium, equal to the rate of PI(4)P synthesis and hydrolysis. Considering PI(4)P level in HeLa cells (56) and that inhibition of the OSBP leads to a two-fold increase in cellular PI(4)P level in one hour (25), we estimated that the OSBP/SAC1ML axis can use up to 660 µM PI(4)P *per* hour to transfer an equivalent amount of sterol, which is compatible with our first scenario (as 3600 x 0.42 μM. s^−1^ = 1500 µM PI(4)P hydrolyzed *per* hour). Finally, these estimations on sterol transfer/PI(4)P hydrolysis rates suggested that OSBP can renew the Golgi sterol pool (400 µM) in one hour, in line with the notion that this protein ensures 30-60 % of ER-to-Golgi sterol transfer (25).

### Sec14p-mediated PI transfer between membranes promotes Osh4p-mediated sterol transfer *via* PI(4)P metabolism

In yeast, Sec14p is a PI/PC exchanger that works with the PI 4-kinase Pik1p to maintain PI(4)P level at the Golgi complex, thereby controlling the secretory function of this organelle. Intriguingly, the activities of Sac1p and Osh4p counteract those of Sec14p and Pik1p (43, 46–48). Because the ER is the source of PI (41), these observations suggest that Sec14p and more generally PI-transfer protein moves PI from this organelle to the *trans*-Golgi and PM to support PI-to-PI(4)P conversion and possibly ORP-mediated lipid exchange. To test this idea, we examined whether Sec14p could transfer PI from ER to Golgi mimetic membranes and thereby promote PI(4)P synthesis by PI4K. L_E_ liposomes composed of DOPC/soy PI/Rhod-PE (78/20/2) were incubated with an equal amount of L_G_ liposomes devoid of PI and decorated with PI4K **(Fig. 6A)**. Soy PI was selected for these assays because, comparatively to liver PI, it is more similar to yeast PI species (3), and properly recognized by Sec14p (57). NBD-PH_FAPP_ was mixed with these liposomes; then Sec14p (200 nM) and ATP (100 µM) were added simultaneously. This elicited a high increase in fluorescence within one hour, suggesting that NBD-PH_FAPP_ associated with L_G_ membranes, in which no Rhod-PE was present to quench the NBD signal. Without Sec14p or ATP, a much lower and no increase were measured, respectively. All these observations suggested that PI was transferred by Sec14p from L_E_ to L_G_ liposomes and converted into PI(4)P by PI4K in the presence of ATP. Confirming this, we observed that adding Osh4p provoked a decrease in fluorescence, suggesting that neo-synthesized PI(4)P was present in L_G_ membranes and transferred by the protein to L_E_ membranes **(Fig. 6B)**. As expected, mirror experiments, in which Rhod-PE was incorporated in L_G_ instead of L_E_ membranes, provided opposite signals: the fluorescence of NBD-PH_FAPP_ slightly decreased when Sec14p and ATP were added, suggesting that NBD-PH_FAPP_ associated to L_G_ liposomes following PI(4)P synthesis ; then, the subsequent injection of Osh4p provoked an increase of this signal, meaning that the sensor relocated to L_E_ liposomes, following PI(4)P transfer **(Fig. 6C,D)**. Jointly these data suggested that Sec14p could support a kinase-dependent production of PI(4)P in the Golgi membrane through PI delivery and that Osh4p could counteract Sec14p action by removing PI(4)P from this membrane.

**Figure 6.**
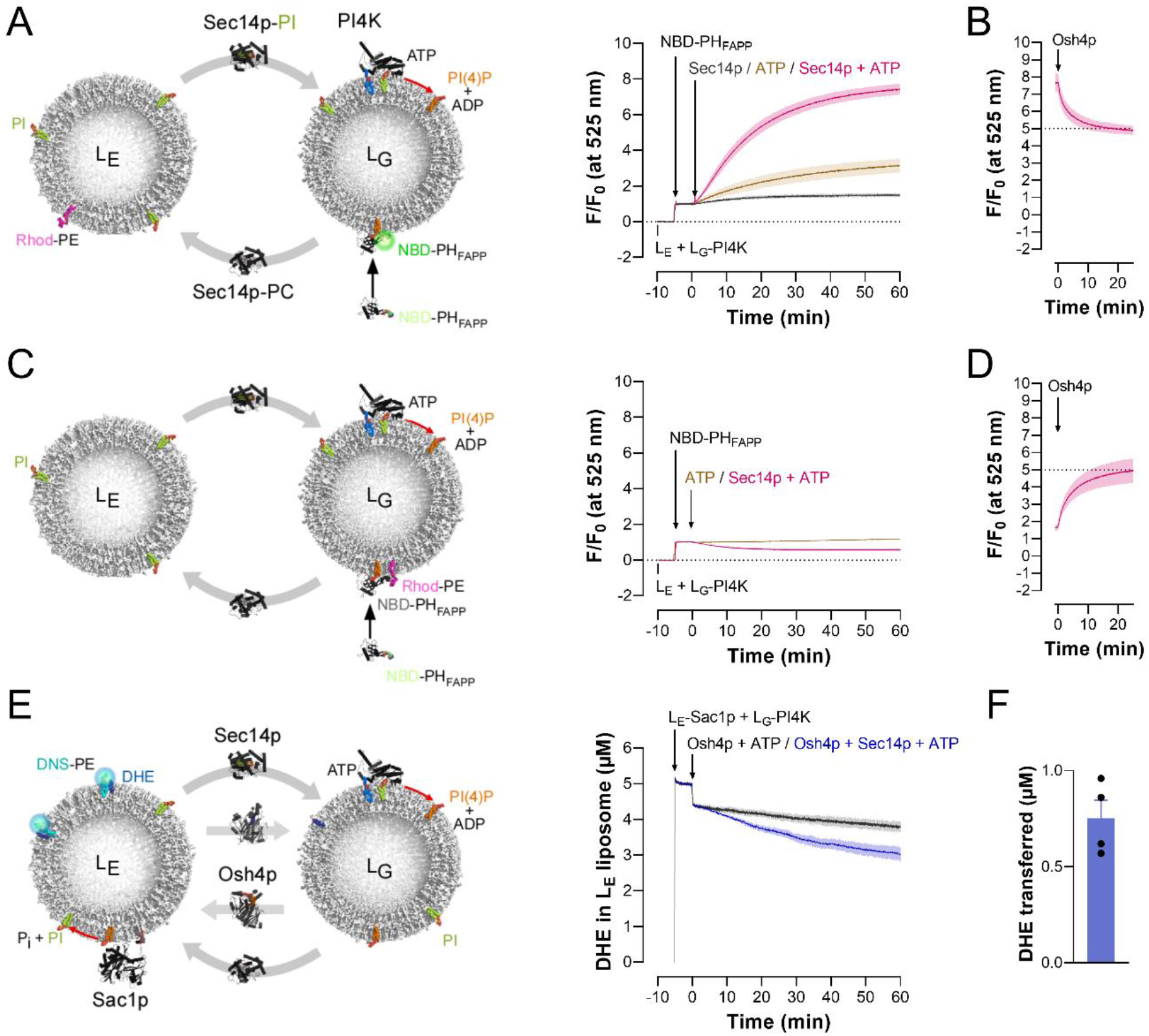
Sec14p-mediated PI transfer supports PI(4)P synthesis and Osh4p activity. (A) L_G_ liposomes (200 µM lipids), composed of DOPC/MPB-PE (97/3) and functionalized with PI4K (PI4K/L=200), were mixed with an equal amount of L_E_ liposomes composed of DOPC/soy PI/Rhod-PE (78/20/2) in HKM7 buffer at 30°C. NBD-PH_FAPP_ (250 nM) was subsequently added. Five minutes after, ATP (100 µM, final concentration) and Sec14p (100 nM) were either injected alone or in association. Fluorescence was measured at 525 nm (λ_ex_ = 460 nm). The signal was normalized to the fluorescence F_0_ measured before the last injection. Mean ± s.e.m. (grey trace, Sec14p only, n = 3; brown trace, ATP only, n = 3; Sec14p pre-mixed with ATP, pink trace, n = 6). (B) At the end of kinetics measured in the presence of Sec14p and ATP, shown in (A), Osh4p (200 nM) was added, provoking a decrease in fluorescence suggestive of a transfer of PI(4)P from L_G_ to L_E_ liposomes. The signal was normalized to F_0_ signal measured in (A). Mean ± s.e.m. (n = 3). (C) Same experiments as in (A) except that 2 mol% Rhod-PE was incorporated in L_E_ liposomes and not in L_G_ liposomes. The signal was normalized to F_0_ signal measured in experiments shown in (A). Mean ± s.e.m. (n = 3). (D) Osh4p (200 nM) was added at the end of kinetics recorded in the presence of Sec14p and ATP, shown in (C). The increase in fluorescence suggested that PI(4)P was transferred from L_G_ to L_E_ liposomes. The signal was normalized to the F_0_ signal measured in (A). Mean ± s.e.m. (n = 3). (E) DHE transfer assays. L_E_ liposomes composed of DOPC/soy PI/DHE/DNS-PE/DOGS-NTA-Ni^2+^ (73/20/2.5/2.5/2), decorated with 100 nM Sac1p[1-522]His_6_, were added to L_G_ liposomes composed of DOPC/MPB-PE/DHE (94.5/3/2.5) and functionalized with PI4K (PI4K/L = 1/200) in HKM7 buffer. The concentration of L_E_ and L_G_ liposomes was identical (200 µM lipids). After five minutes, Osh4p (200 nM) and ATP (100 µM), with or without 200 nM Sec14p, were injected. Mean ± s.e.m. (n = 4). (F) Amount of DHE transferred by Osh4p from L_E_ to L_G_ liposomes within 1 hour, in the presence of ATP and Sec14p. Data are represented as mean ± s.e.m. (error bars; n = 4).

Finally, we examined whether Sec14p, by providing PI to PI4K, and thereby supporting PI(4)P synthesis, could promote Osh4p-mediated sterol transfer. To this end, we prepared L_E_ liposomes composed of DOPC/soy PI/DHE/DNS-PE/DOGS-NTA-Ni^2+^ (73/20/2.5/2.5/2), functionalized with Sac1p[1–522]His_6_, and L_G_ liposomes containing 2.5 mol% DHE but no PI, and covered with PI4K **(Fig. 6E)**. Adding Osh4p and ATP did not provoke any noticeable DHE transfer from L_E_ to L_G_ liposomes, suggesting that no PI(4)P was generated to support Osh4p activity. On the contrary, the conjoint addition of Osh4p, Sec14p and ATP resulted in sterol transfer (**(Fig. 6E, F,** 0.75 ± 0.09 µM DHE, n = 4). These data showed that a PI-transfer protein can support the lipid transfer activity of an ORP by promoting PI(4)P synthesis.

## Discussion

How lipid transfer processes are coupled to lipid metabolism is poorly described quantitatively. In particular it is unclear to what extent PI(4)P metabolism drives ORP-mediated lipid transfer and thereby contributes to creating lipid asymmetries throughout the cell (17). To get more quantitative insights, we analyzed *in vitro* how a PI(4)P gradient generated by a PI 4-kinase and Sac1 between two membranes can drive sterol/PI(4)P exchange. Moreover, we examined whether a PI-transfer process can assist sterol/PI(4)P exchange by promoting PI(4)P synthesis.

Supporting our initial model of sterol/PI(4)P exchange (19), various studies provided clues that a PI(4)P gradient at the ER/Golgi, ER/PM, ER/endosome and ER/lysosome interface was able to pilot vectorial sterol or PS transfer by ORPs (20, 22–26, 28, 58). Notably, it has been shown that their transfer activity could be impacted by the inactivation of Sac1 (22, 24). Alternatively it has been shown that ORPs defective in recognizing PI(4)P were unable to transport their second lipid ligand (20, 22–24, 28). Here, using a controlled *in vitro* system, with chemically-defined liposomes and recombinant proteins, we fully demonstrated that PI(4)P metabolism can drive the vectorial transfer of lipids by an ORP and the formation of a lipid gradient between membranes.

First we found that under initial conditions where sterol was evenly distributed between two membranes, Osh4p could enrich one membrane with sterol, as soon as PI(4)P was synthesized in this membrane, at the expense of the other one. Second, we found that the amplitude of the sterol gradient generated by Osh4p was higher in the presence of Sac1p, which hydrolyzes PI(4)P and maintains the PI(4)P gradient generated by the PI 4-kinase and dissipated by sterol/PI(4)P exchange. This last observation is in line with our previous studies using a membrane system with a pre-existing PI(4)P gradient (21). Finally, we showed that ATP consumption was needed for the creation of a lipid gradient between membranes, demonstrating that interorganelle lipid transport also needs energy, similar to the creation of lipid asymmetry across membranes by ATP-dependent flippases and floppases (59).

From a quantitative point-of-view, we observed that during the creation of a sterol gradient, the net transfer of one sterol molecule required the synthesis of more than one molecule of PI(4)P. This seems at odd with our previous study showing that the creation of a sterol gradient by Osh4p can occur by the one-for-one exchange of sterol for PI(4)P (21). However, these results were obtained under conditions where Osh4p could immediately use a pre-existing PI(4)P gradient to generate a sterol gradient. Then, the sterol gradient was quickly dissipated by a re-equilibration of sterol between membranes (21). Here, we reconstituted a more realistic system where PI(4)P was continually synthesized and hydrolyzed. PI(4)P in our experiments builds up slowly both in liposomes membrane and in complexes with Osh4p, which is why transporting 1 sterol molecule requires to produce more than 1 molecule of PI(4)P. In addition, because PI(4)P synthesis and DHE transfer occur slowly in our system, the spontaneous transfer of sterol between liposomes in stirring condition is non-negligible and counteracts the creation of a sterol gradient. These two factors explains well why there is not a one-to-one ratio between the amount of synthesized PI(4)P and transferred sterol in our system.

Considering these observations and parameters such as PI(4)P synthesis and hydrolysis rates, sterol/PI(4)P exchange rate as well as lipid and protein concentration, we built a kinetic model in good agreement with our *in vitro* data. Our modeling approach *in vitro* allowed us to estimate all kinetic parameters (the rate of lipid exchange associated to Osh4p, *r*, the rate of dephosphorylation due to Sac1, *k*_*sac*_ and the rate of PI(4)P production in L_G_ liposomes, Π); it elucidates the speed up of transport thanks to production of PI(4)P in Golgi mimetic membrane and dephosphorylation in ER mimetic membrane and finally it predicts that spontaneous transfer occurs *in vitro*. Interestingly, this model indicates that sterol can be moved by Osh4p from a membrane to a second one as long as the sterol/PI(4)P molar ratio is lower in the second membrane than in the first. Presumably, a similar principle drives vectorial PS transfer by PS/PI(4)P exchange. Therefore, that the total amount of PI(4)P is much lower than that of sterol or PS in cells is not an obstacle *per se* for the vectorial transfer of these lipids between organelles. This is in fact exactly what we observed *in vitro*: Osh4p is perfectly able to establish a vectorial flux of sterol as soon as the synthesis of a few molecules of PI(4)P starts in face of a high amount of sterol. This observation also suggests that as long as the PI(4)P concentration gradient is steeper than that of sterol and PS at the interface between the ER and other organelles, which is indeed the case considering the intracellular distribution of these three lipids (6–12, 60), a vectorial transfer of sterol and PS by ORPs proteins can be guaranteed.

To go further, we estimated how much sterol could be transferred from ER to Golgi membrane under the control of PI(4)P metabolism in yeast and HeLa cells, using our model. Estimations were done by combining parameters derived from *in vitro* data (*i.e.,* intrinsic activity of Osh4p and Sac1p) and others inferred from data on the geometry and lipid levels in the ER and Golgi apparatus and the abundance of Osh4p or OSBP and Sac1 in these two sorts of cell. We conclude that the amount of Golgi PI(4) and PI(4)P turn-over are compatible with the maintenance of sterol level in the Golgi membrane by Osh4p and OSBP. By gathering recent and disseminated data from the literature to feed our model, we also made interesting observations. First, even if sterol is more concentrated in the Golgi than in the ER membranes of HeLa cell, the sterol pools in the ER and Golgi membranes in terms of volumic concentration are predicted to be similar because the ER surface is much higher than that of the Golgi complex (61). Second, our calculations based on a recent lipidomics study (56) suggest that PI(4)P level is comparable to that of sterol in the Golgi apparatus of HeLa cells. In parallel, it has been recently reported that in yeast, sterol is not more concentrated in the Golgi membrane compared to the ER membrane (62). All these factors can facilitate robust sterol/PI(4)P exchange mediated by ORPs at the ER/Golgi interface, and explain previous observations on cellular OSBP transfer activity (25). Our predictions have limitations yet. First, our estimation of sterol and PI(4)P levels in the ER and Golgi membrane likely lacks precision. Notably an accurate estimation of the level of sterol in the Golgi apparatus in yeast is difficult because in this cell type the architecture of this organelle is poorly defined (63) and the number of post-Golgi secretory vesicles is unknown. Second, PI(4)P has also signaling function, meaning that our prediction are likely based on an overestimation of the amount of PI(4)P directly available for exchange with sterol. On the other hand, we did not consider in our model a factor that can enhance the transfer of sterol to the Gogi by ORPs, i.e. the high affinity of sterol for saturated phospholipids and/or sphingolipids enriched in this organelle (21).

It is still unclear how Sec14p translates its ability to exchange PC and PI between membranes into a biological function (55, 57, 64). One hypothesis is that Sec14p uses this property to carry PI from the ER to the Golgi to support PI(4)P production. Alternatively, it has been proposed that Sec14p performs PC/PI exchange only on the Golgi surface to present PI molecule to Pik1p in an optimal configuration for phosphorylation, thereby boosting Pik1p activity (42, 65). Yet there is no direct evidence that these two proteins function and physically interact together (65). Here, our data support the first model, showing that Sec14p can assist the activity of a PI 4-kinase by delivering PI initially contained in another membrane. Likely, this process does not imply any interaction between Sec14p and PI4K as this latter derived from PI4KIIα which is structurally divergent from Pik1p (66). We also established that Osh4p counteracts the action of Sec14p as found in yeast at the Golgi level (46). Importantly, we observed that Sec14p can promote the sterol/PI(4)P exchange activity of Osh4p by supporting PI(4)P synthesis. This is in line with the idea that PI-transfer proteins can support the activity of ORPs, as suggested for PITPβ and OSBP at the interface between the ER and PI(4)P-rich replication organelles in cells infected by viruses (67). We can thus propose that PI-transfer proteins contribute to equilibrating newly-made PI at the ER/Golgi interface to maintain PI 4-kinase activity and ORP-mediated lipid exchange and possibly recycle PI produced by Sac1p-mediated hydrolysis. Future investigations using our *in vitro* system and in cells are needed to test this model.

At the core of our reconstitution assays is a PI 4-kinase that has been re-engineered to be easily attached to liposomes *via* the formation of a covalent link between its palmitoylated motif and a phospholipid (51). Intriguingly, the kinase was highly active only when in stable association with liposomes, likely because it was locally highly concentrated on membrane surface and able to function in a processive mode with a continuous access to PI. This might also result from a lower mobility of the palmitoylated motif and of an adjacent amphipathic helix close to the catalytic site of the kinase, provoked by their insertion in the membrane (51). However, when PI4K was lying on a membrane that was less fluid as it was made of mono-unsaturated instead of di-unsaturated PC, it did not gain efficiency. In contrast, its activity was found to be dependent on the acyl chain composition of its lipid substrate (unsaturated vs polyunsaturated), as other lipid-synthesizing enzymes and notably PI4 5-kinase (68, 69). Whatever, to our knowledge, we established for the first time a PI(4)P synthesis rate for a PI 4-kinase on a membrane. Yet, it is unclear whether this reflects the activity of the native protein. Indeed, the production of PI(4)P is low, maybe because the modified protein lacks a small N-terminal region or does not adopt an optimal orientation at the membrane interface, *via* its attachment to a phospholipid, in contrast to the palmitoylated form of the protein. It will be of interest to develop a more active kinase for future studies on PI(4)P-driven exchange or for designing screening assays to identify drugs against class II PI-4 kinases (70).

To summarize, our study fully demonstrates that PI(4)P synthesis and hydrolysis on distinct membranes can pilot the vectorial transfer of lipids by ORPs between these membranes, and predict that in cells, PI(4)P turn-over is powerful enough to maintain sterol levels at the ER/Golgi interface by sterol/PI(4)P exchange. Eventually, it shows that PI-transfer proteins can assist the activity of ORPs by supporting PI(4)P synthesis.

### Experimental procedures

#### Protein expression, labelling and purification

Osh4p(C98S), Osh4p(C98S/H143A/H144A), NBD-PH_FAPP_, Sac1p[1-522]His_6_ and Sac1p[1-522](C392S)His_6_ were purified as previously described (21, 24, 54). Their concentration was determined by UV spectrometry.

A truncated version of the human PI 4-kinase IIα (Uniprot Q9BTU6, segment 78-453) whose palmitoylation motif ^174^CCPCC^178^ is replaced by a SSPSS sequence has been previously cloned in PGEX-6P-1 to code for a construct (PI4KIIα^SSPSS^ΔC) in fusion to GST (51). The sequence of this construct was mutated to insert a cysteine at position 178 and to substitute three endogenous and solvent-exposed cysteine residues (C124, C183 and C320) by serine, using the Quikchange kit. All the mutations were checked by DNA sequencing. The GST-PI4KIIα^SSPSC^ΔC(C124S/C183S/C320S) construct was expressed in *E. Coli* (BL21-GOLD(DE3)) competent cells (Stratagene) grown in Luria Bertani Broth (LB) medium at 16°C overnight upon induction with 0.3 mM isopropyl β-D-1-thiogalactopyranoside (IPTG) when the optical density of the bacterial suspension, measured at 600 nm (OD_600_), reached a value of 0.6. Bacteria cells were harvested and re-suspended in cold buffer (50 mM Tris, pH 7.4, 150 mM NaCl, 2 mM DTT) supplemented with 1 mM PMSF, 10 µM bestatin, 1 µM pepstatin A and cOmplete EDTA-free protease inhibitor tablets (Roche). Cells were lysed in a Cell Disruptor TS SERIES (Constant Systems Ltd.) and the lysate was centrifuged at 186,000 × g for 90 minutes. Then, the supernatant was applied to Glutathione Sepharose 4B (Cytiva) for 4 hours at 4°C.

The beads were washed with a buffer containing 50 mM Tris, pH 7.4, 50 mM KCl, 20 mM MgCl_2_ and 5 mM DTT buffer, incubated two times with the same buffer supplemented with 5 mM ATP for 5 and 10 minutes. Finally, the beads were washed with 50 mM Tris, pH 7.4, 150 mM NaCl buffer supplemented with 2 mM DTT and collected in a 2 mL-volume tube. The beads were incubated with PreScission Protease (Cytiva) overnight at 4°C under agitation to cleave off the PI 4-kinase from the GST domain. The protein was recovered in the supernatant after several cycles of centrifugation and washing of the beads, concentrated in 50 mM Tris, pH 7.4, 150 mM NaCl buffer, supplemented with 1 mM DTT, and injected onto a XK-16/70 column packed with Sephacryl S-200 HR to be purified by size-exclusion chromatography. The fractions with ∼100 % pure PI 4-kinase were pooled, concentrated and supplemented with 10 % (v/v) pure glycerol (Sigma). Aliquots were prepared, flash-frozen in liquid nitrogen and stored at -80°C. The concentration of the protein was determined by measuring its absorbance at λ = 280 nm (ε = 63,370 M^-1^.cm^-1^).

GST-Sec14p was expressed in *E. Coli* BL21-GOLD(DE3) competent cells at 30°C overnight upon induction with 1 mM IPTG (at OD_600_ = 0.6). Harvested bacteria cells were re-suspended in 50 mM Tris, pH 7.4, 150 mM NaCl buffer containing 2 mM DTT and supplemented with 1 mM PMSF, 10 µM bestatin, 1 µM pepstatin A and cOmplete EDTA-free protease inhibitor tablets. Cells were lysed and the lysate was centrifuged at 186,000 × g for 1 hour. Next, the supernatant was applied to Glutathione Sepharose 4B for 4 hours at 4°C. The beads were washed three times with protease inhibitors-free buffer and incubated with thrombin at 4°C for 16 hours to cleave off the protein from the GST domain. The protein was recovered in the supernatant after several cycles of centrifugation and washing of the beads, concentrated and purified on a XK-16/70 column packed with Sephacryl S-200 HR. The fractions with ∼100 % pure Sec14p were pooled, concentrated and supplemented with 10 % (v/v) pure glycerol. Aliquots were prepared, flash-frozen in liquid nitrogen and stored at -80°C. The concentration of the protein was determined by measuring its absorbance at λ = 280 nm (ε = 43,570 M^-1^.cm^-1^).

#### Lipids

DOPC (1,2-dioleoyl-*sn*-glycero-3-phosphocholine), liver PI (L-α-phosphatidylinositol, bovine), soy PI, brain PI(4)P (L-α-phosphatidylinositol 4-phosphate), NBD-PE (1,2-dioleoyl-*sn*-glycero-3-phosphoethanolamine-N-(7-nitro-2-1,3-benzoxadiazol-4-yl)), Rhod-PE (1,2-dipalmitoyl-*sn*-glycero-3-phosphoethanolamine-N-(lissamine rhodamine B sulfonyl)), DOGS-NTA-Ni^2+^ (1,2-dioleoyl-*sn*-glycero-3-[(N-(5-amino-1-carboxypentyl)iminodiacetic acid)succinyl] (nickel salt)), MPB-PE (1,2-dioleoyl-*sn*-glycero-3-phosphoethanolamine-N-[4-(p-maleimidophenyl) butyramide]) and DNS-PE (1,2-dioleoyl-*sn*-glycero-3-phosphoethanolamine-N-(5-dimethylamino-1-naphthalenesulfonyl)) were purchased from Avanti Polar Lipids. Dehydroergosterol (DHE) were from Sigma Aldrich. The concentration of DHE in stock solution in methanol was determined by UV-spectroscopy using an extinction coefficient of 13,000 M^-1^.cm^-1^.

#### Liposomes preparation

Lipids stored in stock solutions in CHCl_3_ or CHCl_3_/methanol were mixed at the desired molar ratio. The solvent was removed in a rotary evaporator under vacuum. If the flask contained a mixture with PI(4)P or DOGS-NTA-Ni^2+^, it was pre-warmed at 33°C for 5 minutes prior to creating vacuum. The lipid film was hydrated in 50 mM HEPES, pH 7.4, 120 mM K-Acetate (HK) buffer to obtain a suspension of multi-lamellar vesicles. The multi-lamellar vesicles suspensions were frozen and thawed five times (except for MBP-PE containing liposomes) and then extruded through polycarbonate filters of 0.2 µm pore size using a mini-extruder (Avanti Polar Lipids). Liposomes were stored at 4°C and in the dark when containing fluorescent lipids and used within 2 days.

#### Preparation of liposome functionalized with PI4K

The day of the experiment, a volume of 100 µL from a stock solution of PI4K was applied onto an Zeba column equilibrated with freshly degassed HK buffer, according to manufacturer’s indications, to remove DTT from the protein. The concentration of the eluted protein was determined by UV-spectroscopy. Protein at a final concentration of 1, 2 or 4 µM was then mixed with liposomes (800 µM lipids) composed of DOPC/PI/MPB-PE (87/10/3 mol/mol) for 1 hour at 25°C under agitation in HK buffer. The attachment of PI4K to liposomes was stopped by adding 1 mM DTT.

#### Flotation assay

The association of PI4K protein with membrane was measured by incubating the DTT-free form of the protein (4 µM) with liposomes (800 µM lipids) only made of DOPC, or additionally doped with 3 mol% MPB-PE, 10 mol% liver PI or both lipids, in 150 µL of HK buffer at 25°C for 60 minutes under agitation. Alternatively, to test the respective capacity of PI4K and Sac1p[1-522]-His_6_ to associate with L_E_ and L_G_ membranes, a suspension of L_G_ liposome (520 µM lipids) composed of DOPC/soyPI/DHE/MBP-PE/Rhod-PE (84.5/10/2.5/3/0.1) and functionalized at PI4K/L = 1/200 (1.3 µM kinase, final concentration) were incubated with a suspension of L_E_ liposome (DOPC/DHE/NBD-PE, 97.5/2.5/0.2, 520 µM total lipids), doped or not with DOGS-NTA-Ni²^+^ (at the expense of DOPC) and Sac1p[1-522]-His_6_ (0.5 µM final concentration) in 150 µL of HK buffer for 10 minutes at 25°C under agitation. In each case, the suspension was then adjusted to 28 % (w/w) sucrose by mixing 100 µL of a 60 % (w/w) sucrose solution in HK buffer and overlaid with 200 µL of HK buffer containing 24 % (w/w) sucrose and 50 µL of sucrose-free HK buffer. The sample was centrifuged at 240,000 × g in a swing rotor (TLS 55 Beckmann) for 1 hour. The bottom (250 µL), middle (150 µL) and top (100 µL) fractions were collected. The bottom and top fractions were analysed by SDS-PAGE by direct fluorescence and after staining with SYPRO Orange, using a FUSION FX fluorescence imaging system. For experiments with PI4K and Sac1p, the relative amount of L_E_ and L_G_ liposomes in these fractions, diluted 10-fold in HK buffer, was determined by measuring the fluorescence of NBD-PE (λ_ex_ = 460 nm, λ_em_ = 540 nm) and Rhod-PE (λ_ex_ = 560 nm, λ_em_ = 590 nm) in a 96-well black plate using a Tecan Infinite M1000.

#### Differential flotation assay

“Heavy” L_G_ liposomes (800 µM) composed of DOPC/soyPI/DHE/MBP-PE/Rhod-PE (84.5/10/2.5/3/0.2) and loaded with 50 mM HEPES, pH 7.4, 220 mM sucrose buffer, were functionalized with PI4K (at PI4K/L = 1/200). “Light” L_E_ liposomes (1.3 mM lipids) composed of DOPC/DHE/NBD-PE (97.5/2.5/0.2), doped or not with DOGS-NTA-Ni^2+^ (2 mol% at the expense of DOPC), were prepared in HK buffer and mixed with Sac1p[1-522]His_6_ (0.7 µM) for 10 minutes at 25°C under agitation. Then, 78.125 µL of L_G_ liposome suspension were mixed with 47 µL of the L_E_ liposome suspension in a final volume of 125 µL HK buffer for 10 minutes in a centrifugation tube under constant shaking at 25°C. This suspension was adjusted to 7.5 % sucrose by mixing 75 µL of a 20 % (w/w) sucrose HK buffer then overlaid with 200 µL of 5 % (w/w) sucrose HK buffer, with 200 µL of a 2.5 % (w/w) sucrose HK buffer and finally with 200 µL of sucrose-free HK buffer. The suspension was centrifuged at 240,000 × g in a swing rotor for 1 hour. The bottom (200 µL), middle 1 (200 µL), middle 2 (200 µL) and top (200 µL) fractions were collected. The amount of PI4K and Sac1p[1-522]His_6_ was quantified in each fraction by SDS-PAGE whereas the relative amount of L_E_ and L_G_ liposomes in these fractions was determined by fluorescence measurements.

#### Purification of liposomes functionalized with PI4K

DTT-free PI4K (2 µM) was mixed with liposomes (800 µM lipids) composed of DOPC/PI/MPB-PE (87/10/3) for 1 hour at 25°C under agitation in HK buffer. Then the functionalization reaction was stopped by adding 1 mM DTT. For some assays, liposomes were doped with 2.5 mol% DHE. In each case, control assays were systematically performed with liposomes with the same lipid composition but devoid of MPB-PE and mixed with PI4K. Thereafter, a 150-µL volume of the liposome suspension was adjusted to 28 % (w/w) sucrose by mixing 100 µL of a 60 % (w/w) sucrose solution in HK buffer and overlaid with 200 µL of HK buffer containing 24 % (w/w) sucrose and 50 µL of sucrose-free HK buffer. The sample was centrifuged at 240,000 × g in a swing rotor for 1 hour. The bottom (250 µL), middle (150 µL) and top (100 µL) fractions were collected. The top fractions in which liposomes are concentrated (lipid concentration ∼1200 µM) were used for subsequent experiments.

#### Kinetics assays

All kinetics assays were carried out in a Shimadzu RF 5301-PC fluorimeter. The sample (volume 600 µL) was placed in a cylindrical quartz cell, continuously stirred with a small magnetic bar and thermostated. At the indicated time, sample was injected from stock solutions through a guide in the cover of the fluorimeter adapted to Hamilton syringes, such as to not interrupt the fluorescence recording.

#### Measurement of PI4K–mediated PI(4)P synthesis

A suspension of liposome decorated with PI4K or control liposomes (800 or 1200 µM lipids), was diluted in a final volume of 600 µL of HK buffer containing 7 mM MgCl_2_ in a cuvette thermostated at 30°C. The final concentration of liposome was 200 µM lipids. After 5 minutes, NBD-PH_FAPP_ (250 nM final concentration) was added. Five minutes later, ATP (100 µM final concentration) was injected from a stock solution (10 mM). The fluorescence of NBD-PH_FAPP_ was measured at λ_em_ = 525 nm (bandwidth = 5 nm) on excitation at λ = 495 nm (bandwidth = 5 nm) with a time resolution of 1 s. To measure PI(4)P extraction by Osh4p at the end of the PI(4)P synthesis reaction, a 3 µL-volume of a stock solution of the protein (100 µM) was added every 2 minutes to the sample and the NBD fluorescence was measured between each injection.

#### Determination of PI(4)P production from fluorescence-based assay

To determine the amount of PI(4)P produced by the kinase from the fluorescence trace recorded using NBD-PH_FAPP_, the fluorescence of the sensor was measured in the presence of a series of liposomes, each with a composition corresponding to a given level of PI-to-PI(4)P conversion. Liposomes (200 µM lipids) made of DOPC or POPC and containing different molar ratio of liver PI and PI(4)P (10/0, 9.9/0.1, 9.75/0.25, 9.5/0.5, 9.25/0.75, 9/1, 7.5/2.5, 5/5, 2.5/7.5 and 0/10) were mixed with 250 nM NBD-PH_FAPP_ in 600 µL of HKM7 buffer at 30°C. NBD fluorescence was recorded for 4 minutes at λ_em_ = 525 nm (bandwidth = 5 nm) with λ_ex_ = 495 nm (bandwidth = 5 nm). The intensity values were corrected for the blank (signal of liposome only) and plotted as a function of PI(4)P concentration (from 0 to 10 µM). Data points were fitted using the equation F = F_0_ + F_max_ × ([PI(4)P]/([PI(4)P] + K_D_) where F is the fluorescence of the sensor for a given PI(4)P concentration, F_0_ is the basal fluorescence of the sensor in the presence of liposome devoid of PI(4)P, F_max_ is the fluorescence of the sensor when completely bound to liposome and K_D_ is the dissociation constant of the sensor-membrane binding reaction. With liposomes made of DOPC, the fitting procedure gave F_0_ = 24.2, F_max_ = 78.23 and K_D_ = 0.742 whereas with liposomes made of POPC, the fitting procedure gave F_0_ = 17.15, F_max_ = 93.65 and K_D_ = 1.112. The fluorescence traces were corrected for photobleaching, which was determined by measuring for 1 hour the slow decrease in fluorescence of the NBD-PH_FAPP_ mixed with liposomes composed of DOPC/liver PI/PI(4)P (90/9/1). The rate of PI(4)P synthesis (K_SYN_) was then determined by fitting the data points (F) recorded for 1 hour after ATP injection using the equation F = F_max_ × ([PI(4)P]/ [PI(4)P] + K_D_) with [PI(4)P]=10 × (1-exp(-kt)) ; the conversion of PI into PI(4)P was considered to be a pseudo first-order reaction (as [ATP] >> [PI]). The k value was determined from each fluorescence trace and the corresponding kinetic of PI(4)P synthesis was calculated over a period of 1 hour.

#### DHE transport assay

Liposomes (termed L_G_, 800 µM lipids), composed of 84.5 mol% DOPC (or POPC), 10 mol% PI (soy or liver), 3 mol% MPB-PE and 2.5 mol% DHE, were functionalized with DTT-free PI4K (1, 2 or 4 µM final concentration) for 1 hour at 25°C under agitation. The reaction was stopped by adding 1 mM DTT. Thereafter, a 150-µL volume of the L_G_ liposome suspension was mixed with 420 µL of HK buffer supplemented with 10 mM MgCl_2_ in a cuvette thermostated at 30°C under agitation. Alternatively, a 100-µL volume of purified L_G_ liposomes was mixed with 470 µL of HK buffer supplemented with 8.9 mM MgCl_2_. Five minutes after, a 33-µL volume of a suspension of liposome composed of DOPC/DHE/DNS-PE/DOGS-NTA-Ni^2+^ (93/2.5/2.5/2 mol/mol, L_E_ liposomes, 4 mM lipids), and pre-mixed with Sac1p[1-522]His_6_ or Sac1p[1-522](C392S)His_6_ (1.8 µM), were injected. Then, 5 minutes after, 12 µL of Osh4p(C98S) (stock solution 10 µM), or 12 µL of a mixture of Osh4p with ATP (5 mM), were injected. DHE is quickly equilibrated between the inner and outer leaflet of liposomes (t_1/2_ of transbilayer movement <1 minute (71)) implying that L_E_ and L_G_ liposomes contained each of pool of 5 µM that is accessible to Osh4p. Lipid transport was followed by measuring the dansyl signal at λ_em_ = 525 nm (bandwidth 10 nm) on DHE excitation at λ = 310 nm (bandwidth 1.5 nm). At the end of the kinetics, a final quantity of 10 mM methyl-β-cyclodextrin was added to fully extract DHE from liposomes. The amount of DHE (in µM) transferred from L_E_ to L_G_ liposomes corresponds to 5 × ((F-F_0_)/(F_max_-F_0_)). F_max_ is the signal before Osh4p injection and F_0_ is the signal measured at the end of the kinetics after the addition of methyl-β-cyclodextrin.

#### Measurement of PI(4)P transport coupled to PI(4)P synthesis

Liposomes (termed L_G_, 800 µM lipids) composed of DOPC/liver PI/ MPB-PE/Rhod-PE (85/10/3/2) were incubated with DTT-free PI4K (4 µM final concentration) for 1 hour at 25°C under agitation. Liposome functionalization was stopped by adding 1 mM DTT. Thereafter, 150 µL of the L_G_ liposome suspension were mixed with 420 µL of HK buffer supplemented with 10 mM MgCl_2_ in a cuvette thermostated at 30°C under agitation. After 4 minutes, 30 µL of a suspension of DOPC liposomes (L_E_, 4 mM lipids) were injected. One minute after, NBD-PH_FAPP_ (250 nM) was added to the liposome suspension. Finally, five minutes after, 12 µL of a stock solution of ATP (10 mM), or Osh4p(C98S) (10 µM), or a mix of both was injected. In some experiments, L_E_ and/or L_G_ liposomes were doped with 2.5 mol% DHE at the expense of DOPC. The fluorescence was measured at λ_em_ = 525 nm (bandwidth 10 nm) on excitation at λ_ex_ = 460 nm (bandwidth 1.5 nm).

#### Measurement of PI(4)P synthesis coupled to PI transport

Liposomes (L_G_, 800 µM lipids) composed of DOPC/MPB-PE (97/3) were incubated with DTT-free PI4K (2 µM final concentration) for 1 hour at 25°C. The reaction was stopped by adding 1 mM DTT. Thereafter, a 150-µL volume of this liposome suspension was mixed with 30 µL of a suspension of liposomes composed of DOPC/soy PI/Rhod-PE (78/20/2 mol/mol, L_E_ liposome, 4 mM lipids) and 420 µL of HK buffer supplemented with 10 mM MgCl_2_ in a cuvette thermostated at 30°C. Then, NBD-PH_FAPP_ (250 nM) was added to the liposome suspension. Five minutes after, 100 µM ATP or 200 nm Sec14p, or a mix of both was injected. To perform mirror experiments, Rhod-PE was incorporated in L_E_ and not in L_G_ liposomes. The fluorescence was measured at λ_em_ = 525 nm (bandwidth 5 nm) on excitation at λ_ex_ = 460 nm (bandwidth 5 nm) under constant stirring.

#### PI(4)P-to-DHE exchange assay

We prepared Osh4p(C98S) in complex with PI(4)P as described previously (19) and measured PI(4)P-to-DHE exchange by measuring the quenching of the intrinsic fluorescence of the protein when loading DHE. The sample initially contained 0.5 µM Osh4p(C98S) in HKM buffer at 30°C under continual stirring. Two minutes after, liposomes only made of DOPC or additionally containing 0.1, 0.5, 1, 2 or 5 mol% DHE (500 μM total lipids) were added into the sample from concentrated stock suspension (8 mM lipids). The fluorescence was measured at λ_em_ = 340 nm (bandwidth 5 nm) on excitation at λ_ex_ = 286 nm (bandwidth 1.5 nm).

#### PI 4-kinase assays

PI4K activity was determined by measuring the incorporation of the radioactivity from [γ-^32^P]ATP into PI. DTT-free PI4K (4 µM) was mixed with DOPC liposomes (800 µM lipids) containing 10 mol% PI and either 0 or 3 mol% MPB-PE at 25°C under agitation. After 1 hour, DTT (1 mM) was added to stop the reaction. Then, two series of 1.5-mL tubes containing 50 µL of each liposome preparation, diluted in a final volume of 200 µL of HKM7 buffer, were prepared. In parallel, control tubes only containing DOPC/PI (90/10) liposomes (200 µM) were prepared. Then, 100 µM [γ-^32^P]ATP was added to each tube and the reaction was let to proceed at 30°C under vigorous agitation for 1 hour. Next, 3 × 50 µL from each reaction mix were transferred to glass tubes and mixed with a volume of 3 mL CHCl_3_/CH_3_OH/concentrated HCl, 200:100:0.75 (v/v). The organic phase was then separated from [γ-^32^P]ATP by adding 0.6 ml of 0.6 M HCl solution. Each tube was mixed vigorously and left standing for phase separation. Then, the upper (aqueous) phase was discarded, and 1.5 mL of CHCl_3_:CH_3_OH:0.6 M HCl, (3:48:47, v/v) was added to the lower phase, followed by mixing and phase separation. The upper phase was discarded and the lower phase was transferred to scintillation vials. After solvent evaporation the radioactivity in vials was analyzed by liquid scintillation spectrometry. The amount of synthesized PI(4)P in each reaction mix was determined knowing the radioactivity amount associated to lipids, measured in triplicate, and the amount of radioactivity measured in a 5-µl sample of each reaction mix.

#### ADP-Glo assays

Liposomes functionalized or not with PI4K and purified by flotation (200 µM lipids) were incubated with ultra-pure ATP (100 µM final, V915A, Promega) for 1 hour at 30°C in HKM7 buffer (final volume 100 µL) under constant shaking. Then, the amount of ADP generated by ATP hydrolysis was determined using the ADP-Glo™ Kinase Assay kit (Promega). A 20-µL volume per replicate was mixed with 20 µL of ADP-Glo reagent in a 96-well white plate to stop the kinase reaction and, after 40 minutes at room temperature, to completely remove the unreacted ATP. Then, 40-µL volume of kinase detection reagent were added to transform ADP to ATP and convert this newly-made ATP into light through a luciferase reaction. Once the reaction was completed after one-hour incubation, the sample luminescence was recorded using a Promega™ Glo-max plate reader. The amount of consumed ATP was determined against a standard curve with a range of ATP from 0 to 10 µM.

#### Circular dichroism

The experiments were performed on a Jasco J-815 spectrometer at room temperature with a quartz cell of 0.05 cm path length. Proteins were dialyzed three times against 20 mM Tris, pH 7.4, 120 mM NaF buffer for 30 minutes to remove glycerol contained in protein stocks and to exchange buffer. Each spectrum is the average of ten scans recorded from 185 to 260 nm with a bandwidth of 1 nm, a step size of 0.5 nm and a scan speed of 50 nm.min^-1^. The protein concentration, determined at λ = 280 nm, was in the 5-8 µM range. A control spectrum of buffer was subtracted from each protein spectrum. The percentages of secondary structure of proteins were estimated by analyzing the CD spectra (in the 190-250 nm range) using the BeStSel method provided online (72) and compared to the percentages derived from the analysis of their crystal structure using the DSSP algorithm.

#### Kinetic modeling

A full description of the mathematical model used to analyze the *in vitro* system and provide predictions in cells is given in the Supplementary Methods and Material file.

#### Cellular parameters

##### Size, volume and architecture of a haploid yeast cell

the median volume for a wild-type budding yeast cells is 44 fL (44 µm^3^) (73) corresponding to a sphere of 4.3 µm in diameter. For a yeast of this size, the area of the ER is estimated to be 28 µm^2^ (74). In parallel, considering that a yeast cell contains 30 Golgi compartments, similar to slightly concave disks measuring ∼400 nm in diameter and 50 nm in thickness (63), we estimated that the total area of the Golgi membrane is 34 µm^2^.

##### Protein abundance

there are approx. 18,000 Osh4p copies/cell and 11,000 Sac1p copies/cell (https://www.yeastgenome.org/).

##### Ergosterol content in the ER and Golgi membrane

Considering that the area of the ER is 28 µm^2^ and the cross-sectional area of a phospholipid is approx. 0.6 nm^2^, the ER membrane would be composed of 9.3×10^7^ molecules of phospholipid. According to a recent lipidomic analysis, ergosterol represents 9.7 % of total lipids in the ER membrane (62). Therefore, the ER membrane would contain 9.1×10^6^ molecules of ergosterol. thus composed of 1.13×10^8^ molecules of phospholipid. By the same reasoning, we estimated that the Golgi membrane contains 1.11×10^7^ molecules of ergosterol, given that ergosterol accounts for 9.8 % of lipid in the yeast TGN (6).

##### PI(4)P content in the trans-Golgi membrane

the glycerophospholipid content of a haploid yeast is approx. 28 mg/g CDW(cell dry weight) (5) and thus 3.73 × 10^-5^ mol/g CDW, assuming an average molecular weight of 750 g/mol. PI represents 20 % of glycerophospholipid (5) and PI(4)P represents about 1.5 % of total PI (37). Since the dry weight of a single haploid yeast cell (not in the budding phase) is approx. 50 × 10^−11^ g (75, 76), this implies a PI(4)P content of approx. 3.4×10^6^ molecules/cell. Half of PI(4)P is synthesized in the *trans*-Golgi membrane (35), meaning that the cytosolic leaflet of this compartment contains ∼1.7 × 10^6^ molecules of PI(4)P.

##### Size, volume and architecture of a HeLa cell

the average volume of a HeLa cell is 3 pL (3,000 µm^3^) ; a recent study, providing a complete view of the internal architecture of two individual HeLa cells, revealed that the total area of the ER and Golgi membrane is on average 9150 and 730 µm^2^, respectively (61).

##### Protein abundance

there are ∼591,000 OSBP copies and ∼634,000 SACM1L copies in a HeLa cell (77).

##### Ergosterol content in the ER and Golgi membrane

Considering the area of the ER, its membrane would be composed of 3.05 × 10^10^ molecules of phospholipid. Because cholesterol represent 5 % of total lipids in the ER membrane of mammalian cells (78), there is potentially an ER pool of 1.5 × 10^9^ molecules of cholesterol in this organelle in a HeLa cell. Alternatively, given that OSBP optimally functions as confined at ER-Golgi contact sites, we consider that it might only access to a restricted pool of cholesterol present in the ER sections engaged in these contact sites. Accordingly, considering that the total area of these sections roughly matches the area of the Golgi membrane, we estimate that only 1.2 × 10^8^ molecules of cholesterol are immediately available for OSBP. In parallel, we estimate that the Golgi membrane contains 7.3 ×10^8^ molecules of cholesterol as cholesterol accounts for 30% of lipids in the Golgi complex isolated from Hela cells (79).

##### PI(4)P content in the trans-Golgi membrane

a recent and comprehensive lipidomics analysis of phosphoinositides in HeLa cells indicate there are 2 nmoles PI(4)P/1 × 10^6^ cells (56). Considering that the selective inhibition of OSBP lead to a twofold increase in total PI(4)P level in HeLa cells (20), this suggests that 1.2 × 10^9^ molecules of PI(4)P are available for OBSP at the *trans*-Golgi membrane.

### Statistical analyses

Statistical analyses were performed using Prism (GraphPad). p values < 0.05, < 0.01, and < 0.0001 are identified with 1, 2, and 4 asterisks, respectively. ns: p ≥ 0.05. The number of replicates (n) used for calculating statistics is specified in the figure legends.

## Acknowledgments

We wish to thank Dr. Chang Chen for providing the plasmid coding for GST-PI4KIIα^SSPSS^ΔC construct and Dr. Jeffrey Atkinson for the plasmid coding for GST-Sec14p. This work was supported by a “Défi Modélisation du vivant-2019” grant from CNRS and “ANR-16-CE13-0006” and “ANR-15-IDEX-0” grants from the Agence Nationale de la Recherche. A.S. acknowledges support from the European Research Council (ERC) under the European Union’s Horizon 2020 research and innovation programme (grant agreement No 101002724 RIDING), the Air Force Office of Scientific Research under award number FA8655-20-1-7028, and the National Institutes of Health (NIH) under award number R01DC018789

## Author contributions

G.D. designed and supervised research. N.F., M.M., and G.D. carried out site-directed mutagenesis, produced, purified and labeled all the recombinant proteins of this study. N.F., M.M and G.D. performed the *in vitro* experiments. G.D., N.F and M.M. analyzed the data. N.R. and A.S. developed the mathematical model and obtained the analytic and numerical solutions in vitro and in vivo. G.D and A.S. wrote the manuscript. All of the authors discussed the results and commented on the manuscript.

## Supplementary Figures

**Figure Supplementary 1.**
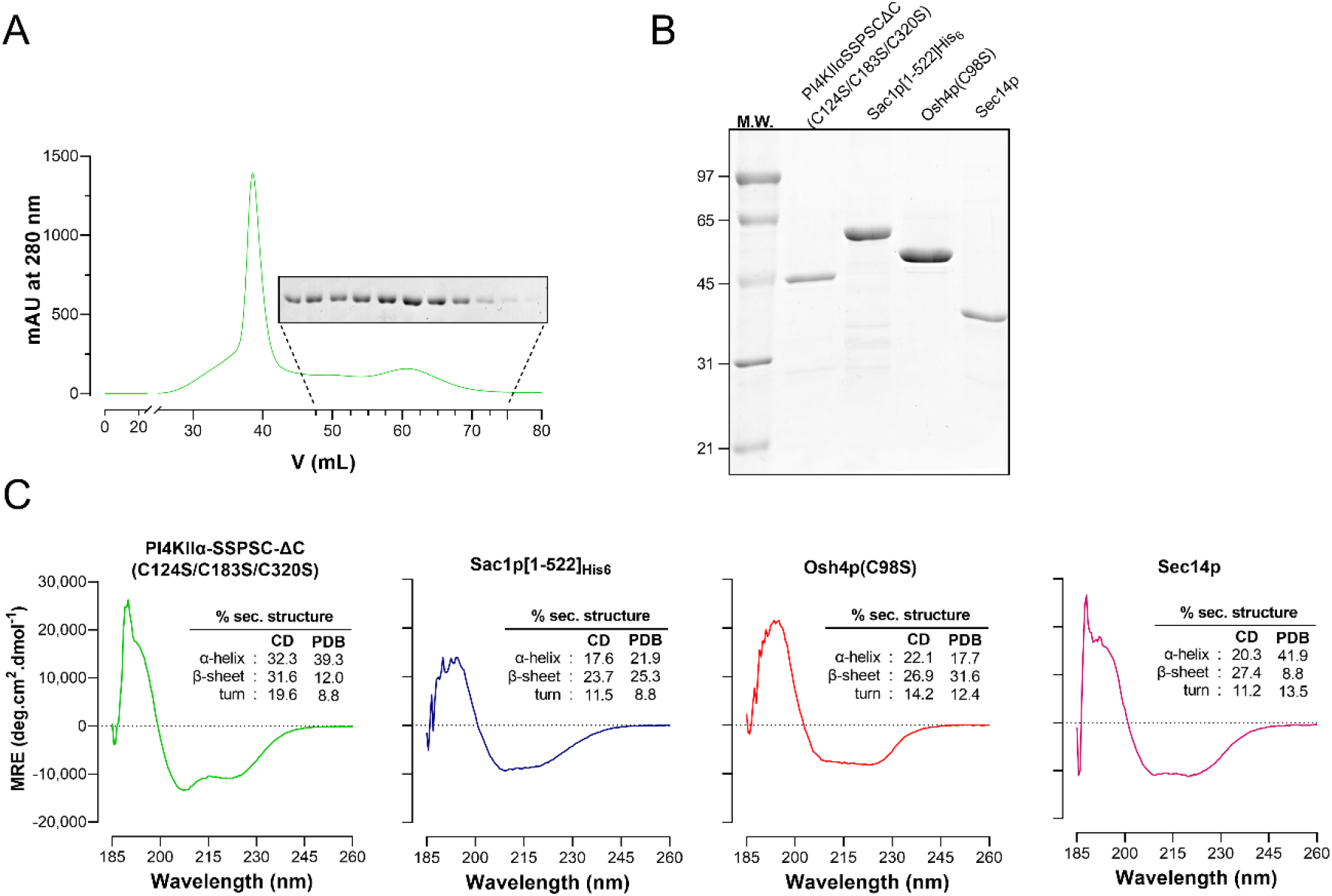
(A) Purification of the PI4KIIα^SSPSC^ΔC(C124S/C183S/C320S) construct. (B) SDS-PAGE analysis of purified PI4KIIα^SSPSC^ΔC(C124S/C183S/C320S), Sac1p[1-522]His_6_, Osh4p(C98S) and Sec14p. The gel was stained with SYPRO Orange to visualize the protein and molecular weight markers. (C) Far-UV CD spectrum of purified PI4KIIα^SSPSC^ΔC(C124S/C183S/C320S), Sac1p[1-522]His_6_, Osh4p(C98S) and Sec14p in 20 mM Tris, pH 7.4, 120 mM NaF buffer. The percentage of α-helix, β-sheet and turn, deriving from the analysis of the CD spectra and from the crystal structures of PI4KIIα^SSPSC^ΔC (PDB ID: 4HNE), Sac1p[1-505] (3LWT), Osh4p (1ZHZ) and Sec14p (1AUA), obtained using the DSSP algorithm, are given. MRE, mean residue ellipticity.

**Figure Supplementary 2.**
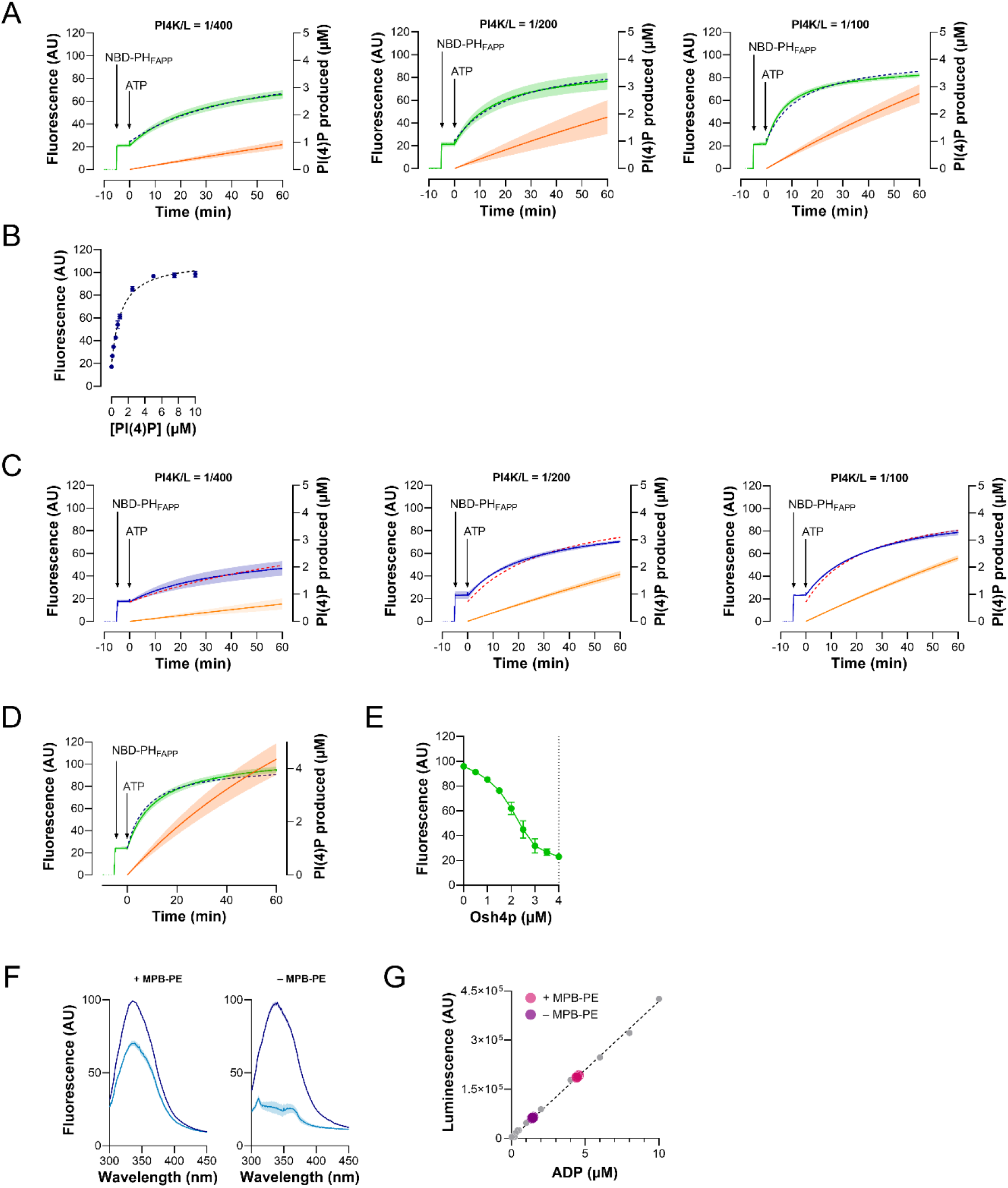
(A) Determination of PI(4)P production. NBD-PH_FAPP_ (250 nM) and ATP (100 µM) were sequentially added to liposomes (200 µM lipids) composed of DOPC/liver PI/MPB-PE (87/10/3) and functionalized with different amounts of PI4K (PI4K/L = 1/100, 1/200 or 1/400). The fluorescence signal was measured at 525 nm (λ_ex_ = 495 nm, green trace) and corrected for photobleaching. The data points were fitted (dotted curve) to determine the rate of PI(4)P synthesis (see Methods). The amount of PI(4)P produced over time by PI4K was then calculated (orange curve). Mean ± s.e.m. (n = 3-4). (B) NBD-PH_FAPP_ (250 nM) was incubated with liposomes (200 µM lipids) made of POPC and containing different amount of PI(4)P at the expense of PI (with PI + PI(4)P representing 10 mol% of total lipids) in HKM7 buffer at 30°C. The fluorescence intensity, measured at 525 nm (λ_ex_ = 495 nm) is reported as a function of the volume concentration of PI(4)P ; the data points were fitted considering that F = F_0_ + F_B_ × [PI(4)P]/([PI(4)P] + K_D_) and a 1:1 stoichiometry. (C) PI(4)P synthesis in POPC membrane. NBD-PH_FAPP_ (250 nM) and ATP (100 µM) were added to liposomes (200 µM lipids), composed of POPC/liver PI/MPB-PE (87/10/3) and covered with PI4K (PI4K/L= 1/100, 1/200 or 1/400). The NBD signal (blue trace), measured at 30°C, was corrected for photobleaching. The data points were fitted to determine the amount of PI(4)P produced over time (orange curve). Mean ± s.e.m. (n = 3-5). (D) PI(4)P synthesis from soy PI. NBD-PH_FAPP_ (250 nM) and ATP (100 µM) were added to liposomes (200 µM lipids) composed of DOPC/soy PI/MPB-PE (87/10/3) and functionalized with PI4K (PI4K/L= 1/200). The fluorescence signal (green trace), measured at 30°C was corrected for photobleaching and used to determine how much PI(4)P was produced within 1 hour (orange curve). Mean ± s.e.m. (n = 7). (E) At the end of kinetics shown in (D), incremental amounts of Osh4p were added to liposomes in which PI(4)P was synthesized. The decrease in fluorescence indicated that Osh4p extracted PI(4)P. A minimal fluorescence value was reached once a total of 4 µM Osh4p was added (dashed lines), suggesting that a corresponding amount of PI(4)P was present in the outer leaflet of liposomes. (F) Prior to kinetics measurements shown in Figure 2F, the intrinsic fluorescence of PI4K associated with PI-containing liposomes (200 µM lipids), doped or not with MPB-PE, without any further purification (blue spectra) and after purification (ligh-blue spectra), was measured (λ_ex_ = 280 nm). Spectra are the mean ± s.e.m. of independent experiments (n = 4). AU, arbitrary units. (G) ATP hydrolysis by PI4K attached to PI-containing liposomes. The panel shows results of a typical experiment. A small sample of liposomes functionalized with PI4K and purified by flotation as shown in Figure 2E, and used in assays shown in Figure 2 F, was diluted 6-fold in 100 µL of HKM7 buffer (200 µM lipids) and mixed with 100 µM ATP. After one-hour incubation at 30°C under constant shaking, the reaction was stopped and the amount of generated ADP was measured (black dots, 5 measurements) by luminescence using an ADP-Glo^TM^ Kinase assay against a standard curve (with a range of ADP from 0 to 10 µM, grey dots). The amount of hydrolysed ATP was also measured for MPB-PE-free liposomes (200 µM lipids, blue dots, 5 measurements), mixed with PI4K and purified by flotation. The data were subtracted from those obtained with liposomes decorated with PI4K, in order to estimate the amount of ATP specifically hydrolysed by the kinase. AU, arbitrary units.

**Figure Supplementary 3.**
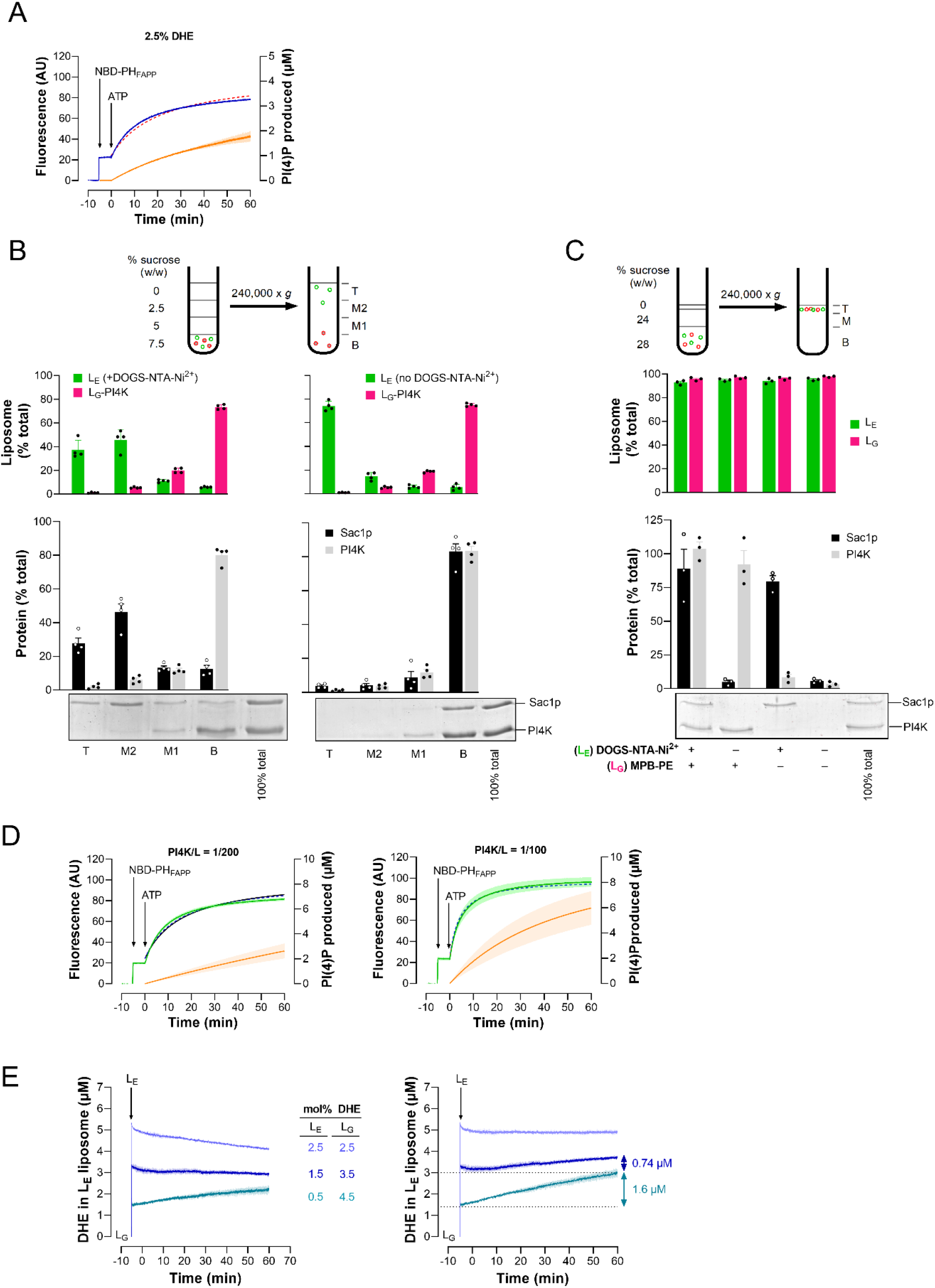
(A) Determination of PI(4)P production in DHE-containing membrane. NBD-PH_FAPP_ (250 nM) and ATP (100 µM) were sequentially added to liposomes (200 µM), composed of DOPC/liver PI/DHE/MPB-PE (84.5/10/2.5/3), and functionalized with PI4K (PI4K/L= 1/100) in HKM7 buffer. The fluorescence signal was measured at 525 nm on excitation at 495 nm (green trace) at 30°C and corrected for photobleaching. The fluorescence data points (green curve) were fitted (dotted curve) to determine the amount of PI(4)P produced over time by PI4K (orange curve). Kinetics traces and derived curves are the mean ± s.e.m. of independent experiments (n = 1-2). (B) Differential flotation assays. Sucrose-loaded L_G_ liposomes composed of DOPC/PI/DHE/MPB-PE/Rhod-PE (84.3/10/2.5/3/0.2, 0.5 mM lipids), functionalized with PI4K (PI4K/L = 200), were mixed with a similar amount of L_E_ liposomes composed of DOPC/DHE/NBD-PE (97.3/2.5/0.2), doped or not with 2 mol% DOGS-NTA-Ni^2+^ (at the expense of DOPC) and with 0.25 µM Sac1p[1-552]His_6_ in HKM7 buffer. The final PI4K concentration is 1.25 µM so that the PI4K/Sac1p concentration ratio is similar to the one in DHE transfer experiments shown in the Figure 3B. The liposomes and proteins were mixed for 10 minutes at 30°C under shaking. After centrifugation on a sucrose gradient, bottom, medium and top fractions were collected. The percentage amount of Sac1p and PI4K in each fraction was determined by SDS-PAGE analysis. The percentage amount of each liposome population in each fraction was evaluated by measuring the fluorescence of NBD-PE and Rhod-PE. Data are represented as mean ± s.e.m. with single data points (n = 4). (C) Flotation assays. L_G_ liposomes composed of DOPC/PI/DHE/Rhod-PE (84.3/10/2.5/0.2, 0.5 mM lipids), doped or not with 3 mol% MPB-PE were incubated for 1 hour with PI4K (PI4K/L = 200) at 30°C, then added to L_E_ liposomes composed of DOPC/DHE/NBD-PE (97/2.5/0.2), containing or not 2 mol% DOGS-NTA-Ni^2+^, pre-mixed for 10 minutes with Sac1p[1-522]His_6_. The final concentration of each liposome population are 500 µM whereas the final concentration of PI4K and Sac1p are 1.3 and 0.5 µM, respectively. After centrifugation, the liposomes were recovered at the top of sucrose cushions and the fraction of PI4K and Sac1p associated to liposomes (lane 1 to 4) was determined by SDS-PAGE using the content of lane 5 (100 % total) as a reference. The percentage amount of L_E_ and L_G_ liposomes in the top fraction was evaluated by measuring the fluorescence of NBD-PE and Rhod-PE in bottom, medium and top fractions. Data are represented as mean ± s.e.m. with single data points (n = 3). (D) PI(4)P synthesis from soy PI in DHE-containing membrane. NBD-PH_FAPP_ (250 nM) and ATP (100 µM) were sequentially added to liposomes (200 µM), composed of DOPC/soy PI/DHE/MPB-PE (84.5/10/2.5/3), and functionalized with PI4K (PI4K/L= 1/200 or 1/400), in HKM7 buffer. The fluorescence data points (green curve) were fitted (dotted curve) to determine the amount of PI(4)P produced by the kinase (orange curve). Kinetics traces and derived curves are the mean ± s.e.m. of independent experiments (n = 3-4). (E) Spontaneous DHE transfer between L_E_ and L_G_ liposomes under conditions where a pre-existing DHE gradient was established between these liposomes. L_G_ liposomes (200 µM) made of DOPC and containing 10 mol% PI, 3 mol% MPB-PE and either 2.5, 3.5 or 4.5 mol% DHE (at the expense of PC) were mixed with L_E_ liposomes (200 µM) containing 2 mol% DOGS-NTA-Ni^2+^, 2.5 mol% DNS-PE and either 2.5, 1.5 or 0.5 mol% DHE. Fluorescence was recorded at 525 nm (λ_ex_ = 310 nm) and normalized in terms of DHE present in L_E_ liposomes. Mean ± s.e.m. (n= 3-4). On the left, the signal was corrected for photobleaching.

**Figure Supplementary 4.**
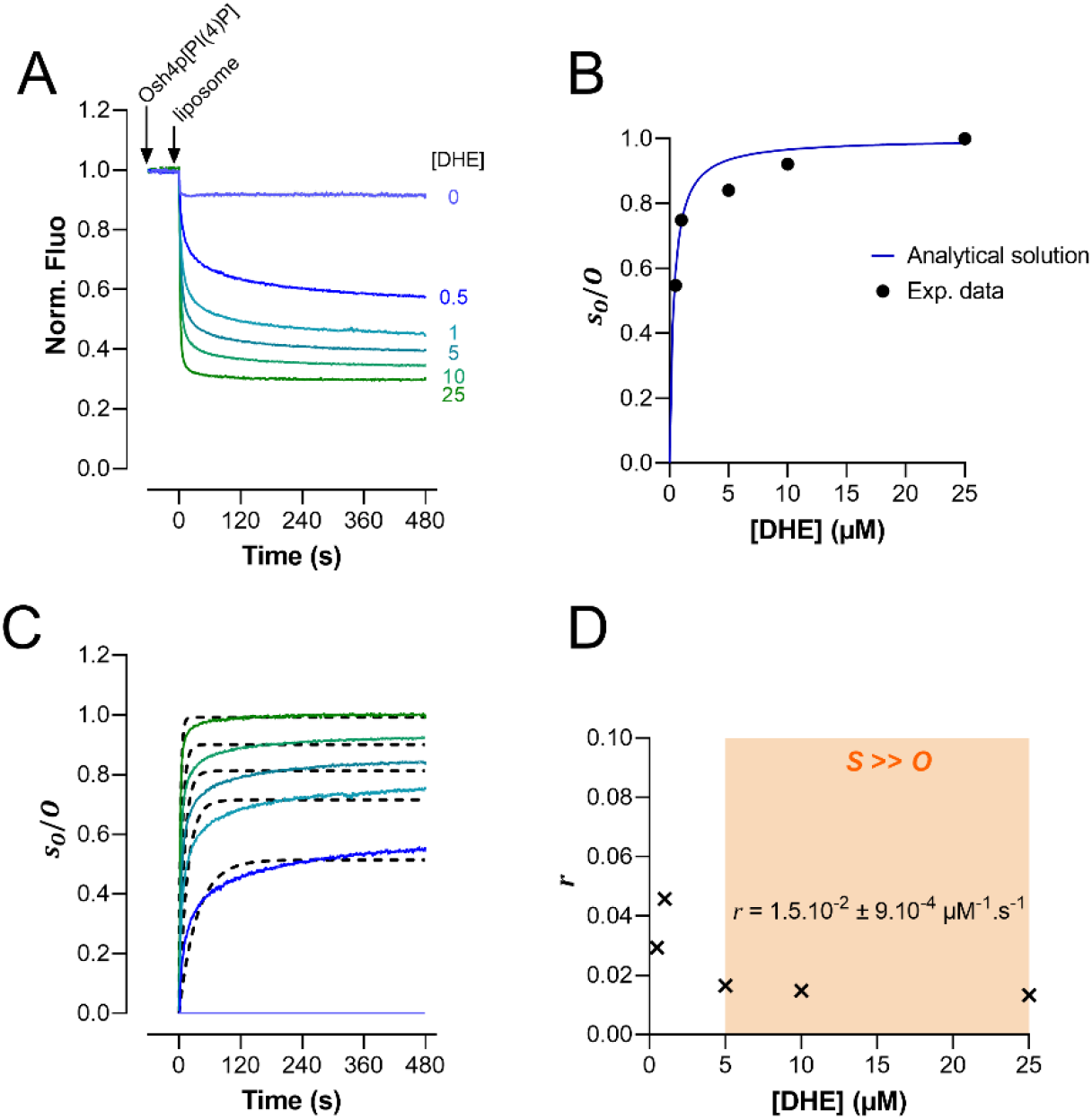
(A) PI(4)P-to-DHE exchange assays. Intrinsic fluorescence of Osh4p(C98S)[PI(4)P] (500 nM) after the addition of liposomes (500 μM total lipids) made of DOPC and containing 0, 0.5, 1, 5, 10, 25 µM DHE. The fluorescence signal was measured at λ_em_ = 340 nm (λ_ex_ = 286 nm). The intensity was normalized to the intensity (F_0_) measured before the last injection. Each trace is the mean ± s.e.m. of independent experiments (n = 3). Next, these curves were normalized to show the kinetics of association of Osh4p with sterol as a function of sterol concentration. The fraction of Osh4p bound to sterols (*s*_*O*_/*O*) was calculated considering that *s*_*O*_/*O* = 1 - (F-F_25_)/(F-F_0_) with F corresponding to the fluorescence signal of Osh4p over time, F_25_ corresponding to the minimum fluorescence of Osh4p measured in the presence of 25 µM DHE and F_0_ corresponding to the fluorescence measured after the addition of Osh4p to DHE-free liposomes. (B) Mean equilibrium value of *s*_*O*_/*O* were calculated from the curves shown in A in the [450-480] seconds interval (black circle) and compared to analytical prediction from our model for protein loading equation (6) Materials and Methods (blue curve). (C) Data points of the kinetics curves for *s*_*O*_/*O* measured at [DHE] = 0.5, 1, 5, 10 and 25 µM (solid lines) has been fitted with the solution (equation (6)) for the model of protein loading, considering that the PI(4)P-to-DHE exchange process follows a second order kinetics. (D) PI(4)P-to-sterol exchange rate (represented as crosses in the right panel) given by the fit shown in C. The mean exchange rate (± s.e.m), established for a regime where DHE is much more abundant than Osh4p (as in Osh4p-mediated DHE transfer assays, orange area) is indicated.

**Figure Supplementary 5.**
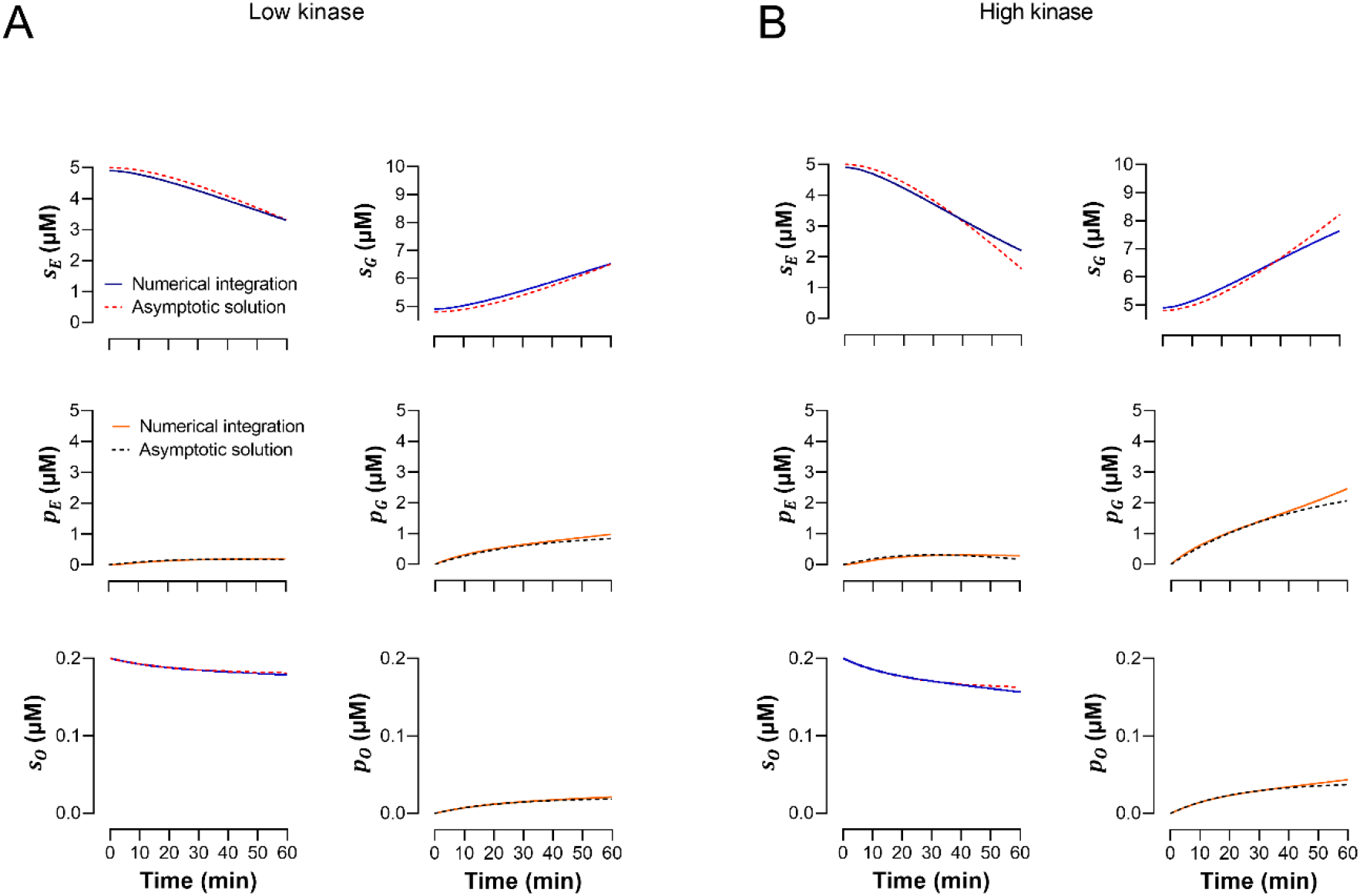
Evolution of the concentration of sterol and PI(4)P in the ER-like membrane (*s*_*E*_, *p*_*E*_) and Golgi-membrane (*s*_*G*_, *p*_*G*_), and of Osh protein in complex with either sterol or PI(4)P (*s*_*o*_, *p*_*o*_) over a period of 1 h, according to the kinetic model equation (9) (Supp. Materials and Methods). The total concentration of Osh protein *O* is 200 nM, the initial concentration of sterol *s*_*E*_ and *s*_*G*_ is equal to 5 µM and the PI(4)P synthesis rate Π is equal to Π = 7 × 10^−4^ μM. s^−1^ (A) and or Π = 1.9 × 10^−3^ µM. s^−1^ (B), similar to conditions of DHE transfer assays (Fig.4A with PI4K/L = 1/200 or PI4K/L =1/100). The values were obtained by integrating numerically the full system (numerical integration, blue or orange line) or by an asymptotic analysis of the full system (asymptotic solution, dashed red or black line).

**Figure Supplementary 6.**
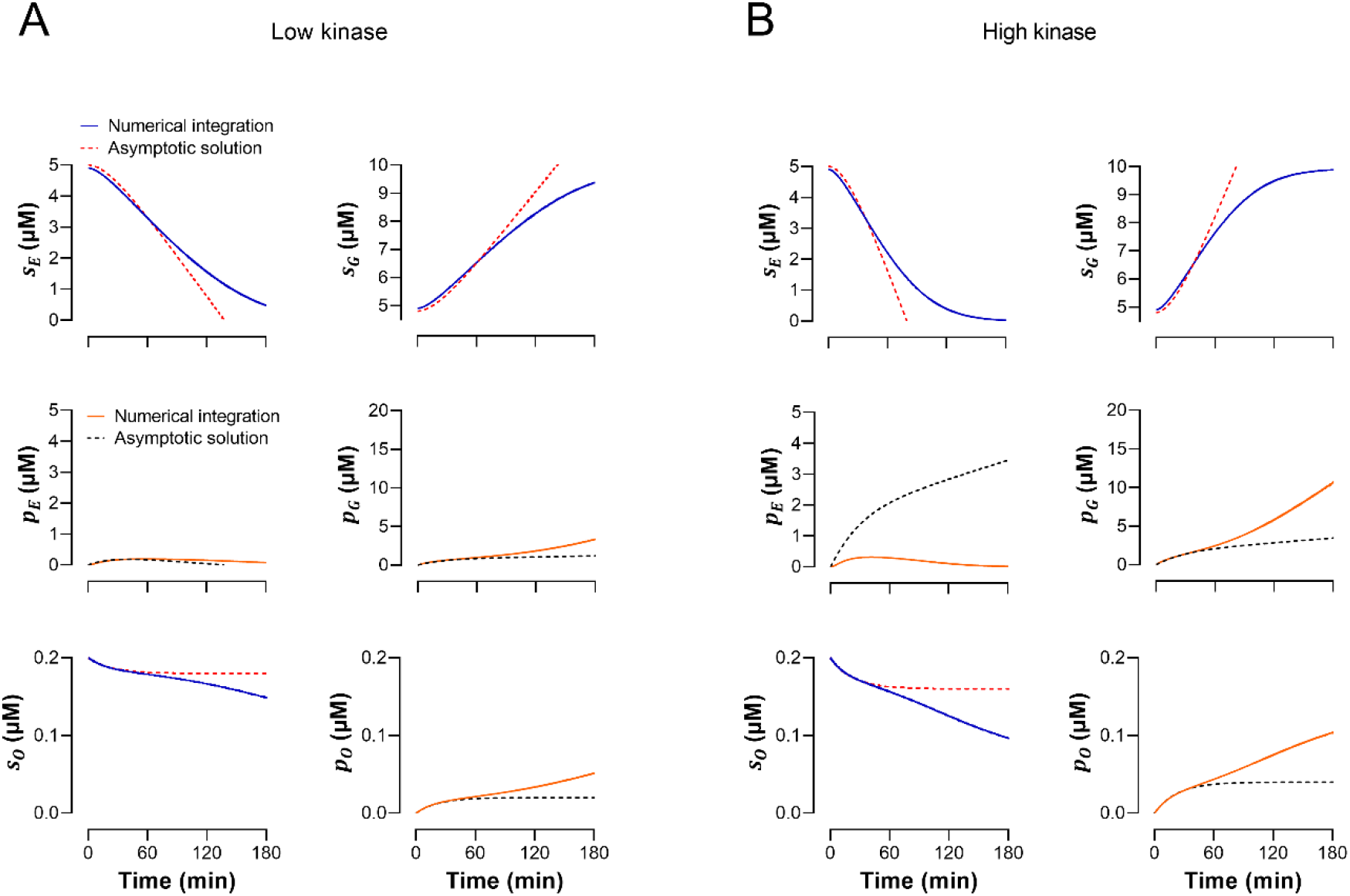
Same as Figure Supplementary 5 but the evolution of the concentration of lipid in ER-like and Golgi-like membranes, and of Osh4p-lipid complexes is shown for a period of 3 hours.

**Table S1.** Estimates for lipid and protein concentration in a yeast and HeLa cell and prediction.

| Concentrations ($\mu\text{M}$) | Yeast | HeLa cells |
| --- | --- | --- |
| $s_E$ | 337 | 67.5 to 844 |
| $s_G$ | 413 | 404 |
| $p_G$ | 62.8 | 664 |
| $O$ | 0.67 (Osh4p) | 0.364 (OSBP) |
| $Sac1$ | 0.41 (Sac1p) | 0.392 (SACM1L) |
| Predicted maximum sterol transport rate $\Pi^*$ ($\mu\text{M} \cdot \text{s}^{-1}$) | 18 | 2.0 to 25 |
| Predicted sterol transport rate $\Pi$ ($\mu\text{M} \cdot \text{s}^{-1}$) | 0.25 | 0.42 to 4 |

## References

1. Holthuis JC & Menon AK (2014) Lipid landscapes and pipelines in membrane homeostasis. Nature 510(7503):48–57.

2. Bigay J & Antonny B (2012) Curvature, lipid packing, and electrostatics of membrane organelles: defining cellular territories in determining specificity. Dev Cell 23(5):886–895.

3. Ejsing CS, et al. (2009) Global analysis of the yeast lipidome by quantitative shotgun mass spectrometry. Proc. Natl. Acad. Sci. U.S.A. 106(7):2136–2141.

4. Sampaio JL, et al. (2011) Membrane lipidome of an epithelial cell line. Proc Natl Acad Sci U S A 108(5):1903–1907.

5. Daum G, et al. (1999) Systematic analysis of yeast strains with possible defects in lipid metabolism. Yeast 15(7):601–614.

6. Klemm RW, et al. (2009) Segregation of sphingolipids and sterols during formation of secretory vesicles at the trans-Golgi network. J Cell Biol 185(4):601–612.

7. Zinser E, et al. (1991) Phospholipid synthesis and lipid composition of subcellular membranes in the unicellular eukaryote Saccharomyces cerevisiae. J. Bacteriol. 173(6):2026–2034.

8. Schneiter R, et al. (1999) Electrospray ionization tandem mass spectrometry (ESI-MS/MS) analysis of the lipid molecular species composition of yeast subcellular membranes reveals acyl chain-based sorting/remodeling of distinct molecular species en route to the plasma membrane. J Cell Biol 146(4):741–754.

9. Andreyev AY, et al. (2010) Subcellular organelle lipidomics in TLR-4-activated macrophages. J Lipid Res 51(9):2785–2797.

10. Vance JE & Steenbergen R (2005) Metabolism and functions of phosphatidylserine. Prog Lipid Res 44(4):207–234.

11. Leventis PA & Grinstein S (2010) The distribution and function of phosphatidylserine in cellular membranes. Annu Rev Biophys 39:407–427.

12. Zinser E, Paltauf F, & Daum G (1993) Sterol composition of yeast organelle membranes and subcellular distribution of enzymes involved in sterol metabolism. J. Bacteriol. 175(10):2853–2858.

13. Mesmin B & Maxfield FR (2009) Intracellular sterol dynamics. Biochim. Biophys. Acta 1791(7):636–645.

14. Levental I, Levental KR, & Heberle FA (2020) Lipid Rafts: Controversies Resolved, Mysteries Remain. Trends Cell Biol. 30(5):341–353.

15. Kay JG & Fairn GD (2019) Distribution, dynamics and functional roles of phosphatidylserine within the cell. Cell Commun Signal 17(1):126.

16. Lenoir G, D’Ambrosio JM, Dieudonne T, & Copic A (2021) Transport Pathways That Contribute to the Cellular Distribution of Phosphatidylserine. Front Cell Dev Biol 9:737907.

17. Drin G (2022) Creating and sensing asymmetric lipid distributions throughout the cell. Emerg. Top. Life Sci.

18. Delfosse V, Bourguet W, & Drin G (2020) Structural and Functional Specialization of OSBP-Related Proteins. Contact 3:2515256420946627.

19. de Saint-Jean M, et al. (2011) Osh4p exchanges sterols for phosphatidylinositol 4-phosphate between lipid bilayers. J. Cell Biol. 195(6):965–978.

20. Mesmin B, et al. (2013) A four-step cycle driven by PI(4)P hydrolysis directs sterol/PI(4)P exchange by the ER-Golgi Tether OSBP. Cell 155(4):830–843.

21. Moser von Filseck J, Vanni S, Mesmin B, Antonny B, & Drin G (2015) A phosphatidylinositol-4-phosphate powered exchange mechanism to create a lipid gradient between membranes. Nat. Commun. 6:6671.

22. Chung J, et al. (2015) INTRACELLULAR TRANSPORT. PI4P/phosphatidylserine countertransport at ORP5- and ORP8-mediated ER-plasma membrane contacts. Science 349(6246):428–432.

23. Encinar Del Dedo J, et al. (2021) Coupled sterol synthesis and transport machineries at ER-endocytic contact sites. J. Cell Biol. 220(10).

24. Moser von Filseck J, et al. (2015) INTRACELLULAR TRANSPORT. Phosphatidylserine transport by ORP/Osh proteins is driven by phosphatidylinositol 4-phosphate. Science 349(6246):432–436.

25. Mesmin B, et al. (2017) Sterol transfer, PI4P consumption, and control of membrane lipid order by endogenous OSBP. EMBO J. 36(21):3156–3174.

26. Radulovic M, et al. (2022) Cholesterol transfer via endoplasmic reticulum contacts mediates lysosome damage repair. Embo j 41(24):e112677.

27. Tan JX & Finkel T (2022) A phosphoinositide signalling pathway mediates rapid lysosomal repair. Nature 609(7928):815–821.

28. Kawasaki A, et al. (2022) PI4P/PS countertransport by ORP10 at ER-endosome membrane contact sites regulates endosome fission. J Cell Biol 221(1).

29. Dong R, et al. (2016) Endosome-ER Contacts Control Actin Nucleation and Retromer Function through VAP-Dependent Regulation of PI4P. Cell 166(2):408–423.

30. Stefan CJ, et al. (2017) Membrane dynamics and organelle biogenesis-lipid pipelines and vesicular carriers. BMC Biol. 15(1):102.

31. Smindak RJ, et al. (2017) Lipid-dependent regulation of exocytosis in S. cerevisiae by OSBP homolog (Osh) 4. J Cell Sci 130(22):3891–3906.

32. Ling Y, Hayano S, & Novick P (2014) Osh4p is needed to reduce the level of phosphatidylinositol-4-phosphate on secretory vesicles as they mature. Mol Biol Cell 25(21):3389–3400.

33. Sohn M, et al. (2016) Lenz-Majewski mutations in PTDSS1 affect phosphatidylinositol 4-phosphate metabolism at ER-PM and ER-Golgi junctions. Proc. Natl. Acad. Sci. U.S.A. 113(16):4314–4319.

34. Sohn M, et al. (2018) PI(4,5)P(2) controls plasma membrane PI4P and PS levels via ORP5/8 recruitment to ER-PM contact sites. J. Cell Biol. 217(5):1797–1813.

35. Audhya A, Foti M, & Emr SD (2000) Distinct roles for the yeast phosphatidylinositol 4-kinases, Stt4p and Pik1p, in secretion, cell growth, and organelle membrane dynamics. *Mol. Biol*. Cell 11(8):2673–2689.

36. Foti M, Audhya A, & Emr SD (2001) Sac1 lipid phosphatase and Stt4 phosphatidylinositol 4-kinase regulate a pool of phosphatidylinositol 4-phosphate that functions in the control of the actin cytoskeleton and vacuole morphology. Mol. Biol. Cell 12(8):2396–2411.

37. Stefan CJ, et al. (2011) Osh proteins regulate phosphoinositide metabolism at ER-plasma membrane contact sites. Cell 144(3):389–401.

38. Stefan CJ, et al. (2017) Membrane dynamics and organelle biogenesis—lipid pipelines and vesicular carriers. BMC Biol. 15(1):102.

39. Pemberton JG, et al. (2020) Defining the subcellular distribution and metabolic channeling of phosphatidylinositol. J Cell Biol 219(3).

40. Zewe JP, et al. (2020) Probing the subcellular distribution of phosphatidylinositol reveals a surprising lack at the plasma membrane. J. Cell Biol. 219(3).

41. Blunsom NJ & Cockcroft S (2020) Phosphatidylinositol synthesis at the endoplasmic reticulum. Biochim. Biophys. Acta Mol.Cell Biol. Lipids 1865(1):158471.

42. Schaaf G, et al. (2008) Functional anatomy of phospholipid binding and regulation of phosphoinositide homeostasis by proteins of the sec14 superfamily. Mol. Cell 29(2):191–206.

43. Hama H, Schnieders EA, Thorner J, Takemoto JY, & DeWald DB (1999) Direct Involvement of Phosphatidylinositol 4-Phosphate in Secretion in the Yeast Saccharomyces cerevisiae *. J. Biol. Chem. 274(48):34294–34300.

44. Grabon A, Bankaitis VA, & McDermott MI (2019) The interface between phosphatidylinositol transfer protein function and phosphoinositide signaling in higher eukaryotes. J. Lipid Res. 60(2):242–268.

45. Lipp NF, Ikhlef S, Milanini J, & Drin G (2020) Lipid Exchangers: Cellular Functions and Mechanistic Links With Phosphoinositide Metabolism. Front. Cell. Dev. Biol. 8:663.

46. Fairn GD, Curwin AJ, Stefan CJ, & McMaster CR (2007) The oxysterol binding protein Kes1p regulates Golgi apparatus phosphatidylinositol-4-phosphate function. Proc. Natl. Acad. Sci. U.S.A. 104(39):15352–15357.

47. Rivas MP, et al. (1999) Pleiotropic alterations in lipid metabolism in yeast sac1 mutants: relationship to “bypass Sec14p” and inositol auxotrophy. Mol. Biol. Cell 10(7):2235–2250.

48. Stock SD, Hama H, DeWald DB, & Takemoto JY (1999) SEC14-dependent secretion in Saccharomyces cerevisiae. Nondependence on sphingolipid synthesis-coupled diacylglycerol production. J. Biol. Chem. 274(19):12979–12983.

49. Roulin PS, et al. (2014) Rhinovirus Uses a Phosphatidylinositol 4-Phosphate/Cholesterol Counter-Current for the Formation of Replication Compartments at the ER-Golgi Interface. Cell Host Microbe 16(5):677–690.

50. Peretti D, Dahan N, Shimoni E, Hirschberg K, & Lev S (2008) Coordinated lipid transfer between the endoplasmic reticulum and the Golgi complex requires the VAP proteins and is essential for Golgi-mediated transport. Mol. Biol. Cell 19(9):3871–3884.

51. Zhou Q, et al. (2014) Molecular insights into the membrane-associated phosphatidylinositol 4-kinase IIα. Nat. Commun. 5:3552.

52. Downing GJ, Kim S, Nakanishi S, Catt KJ, & Balla T (1996) Characterization of a Soluble Adrenal Phosphatidylinositol 4-Kinase Reveals Wortmannin Sensitivity of Type III Phosphatidylinositol Kinases. Biochemistry 35(11):3587–3594.

53. Nakanishi S, Catt KJ, & Balla T (1995) A wortmannin-sensitive phosphatidylinositol 4-kinase that regulates hormone-sensitive pools of inositolphospholipids. Proc. Natl. Acad. Sci. U.S.A. 92(12):5317–5321.

54. Lipp N-F, et al. (2019) An electrostatic switching mechanism to control the lipid transfer activity of Osh6p. Nat. Commun. 10(1):3926.

55. Aitken JF, van Heusden GP, Temkin M, & Dowhan W (1990) The gene encoding the phosphatidylinositol transfer protein is essential for cell growth. J. Biol. Chem. 265(8):4711–4717.

56. Li P & Lämmerhofer M (2021) Isomer Selective Comprehensive Lipidomics Analysis of Phosphoinositides in Biological Samples by Liquid Chromatography with Data Independent Acquisition Tandem Mass Spectrometry. Anal. Chem. 93(27):9583–9592.

57. Sugiura T, et al. (2019) Biophysical Parameters of the Sec14 Phospholipid Exchange Cycle. Biophys. J. 116(1):92–103.

58. Wakana Y, et al. (2021) The ER cholesterol sensor SCAP promotes CARTS biogenesis at ER-Golgi membrane contact sites. J. Cell Biol. 220(1).

59. López-Marqués RL, et al. (2015) Structure and mechanism of ATP-dependent phospholipid transporters. Biochim. Biophys. Acta 1850(3):461–475.

60. Hammond GR, Schiavo G, & Irvine RF (2009) Immunocytochemical techniques reveal multiple, distinct cellular pools of PtdIns4P and PtdIns(4,5)P(2). Biochem. J. 422(1):23–35.

61. Heinrich L, et al. (2021) Whole-cell organelle segmentation in volume electron microscopy. Nature 599(7883):141–146.

62. Reinhard J, et al. (2022) A new technology for isolating organellar membranes provides fingerprints of lipid bilayer stress. bioRxiv:2022.2009.2015.508072.

63. Preuss D, Mulholland J, Franzusoff A, Segev N, & Botstein D (1992) Characterization of the Saccharomyces Golgi complex through the cell cycle by immunoelectron microscopy. Mol. Biol. Cell 3(7):789–803.

64. Szolderits G, Hermetter A, Paltauf F, & Daum G (1989) Membrane properties modulate the activity of a phosphatidylinositol transfer protein from the yeast, Saccharomyces cerevisiae. Biochim. Biophys. Acta 986(2):301–309.

65. Bankaitis VA, et al. (2012) Thoughts on Sec14-like nanoreactors and phosphoinositide signaling. Adv. Biol. Regul. 52(1):115–121.

66. Burke JE (2018) Structural Basis for Regulation of Phosphoinositide Kinases and Their Involvement in Human Disease. Mol. Cell 71(5):653–673.

67. Roulin PS, et al. (2014) Rhinovirus uses a phosphatidylinositol 4-phosphate/cholesterol counter-current for the formation of replication compartments at the ER-Golgi interface. Cell Host Microbe 16(5):677–690.

68. Shulga YV, Anderson RA, Topham MK, & Epand RM (2012) Phosphatidylinositol-4-phosphate 5-kinase isoforms exhibit acyl chain selectivity for both substrate and lipid activator. J. Biol. Chem. 287(43):35953–35963.

69. Barneda D, Cosulich S, Stephens L, & Hawkins P (2019) How is the acyl chain composition of phosphoinositides created and does it matter? Biochem. Soc. Trans. 47(5):1291–1305.

70. Boura E & Nencka R (2015) Phosphatidylinositol 4-kinases: Function, structure, and inhibition. Exp. Cell Res. 337(2):136–145.

71. John K, Kubelt J, Muller P, Wustner D, & Herrmann A (2002) Rapid transbilayer movement of the fluorescent sterol dehydroergosterol in lipid membranes. Biophys. J. 83(3):1525–1534.

72. Micsonai A, et al. (2015) Accurate secondary structure prediction and fold recognition for circular dichroism spectroscopy. Proc. Natl. Acad. Sci. U.S.A. 112(24):E3095–E3103.

73. Jorgensen P, Nishikawa JL, Breitkreutz BJ, & Tyers M (2002) Systematic identification of pathways that couple cell growth and division in yeast. Science 297(5580):395–400.

74. Perktold A, Zechmann B, Daum G, & Zellnig G (2007) Organelle association visualized by three-dimensional ultrastructural imaging of the yeast cell. FEMS Yeast Res. 7(4):629–638.

75. Cuny AP, et al. (2022) High-resolution mass measurements of single budding yeast reveal linear growth segments. Nat. Commun. 13(1):3483.

76. Labedz B, Wanczyk A, & Rajfur Z (2017) Precise mass determination of single cell with cantilever-based microbiosensor system. PLoS One 12(11):e0188388.

77. Itzhak DN, Tyanova S, Cox J, & Borner GHH (2016) Global, quantitative and dynamic mapping of protein subcellular localization. eLife 5:e16950.

78. Radhakrishnan A, Goldstein JL, McDonald JG, & Brown MS (2008) Switch-like control of SREBP-2 transport triggered by small changes in ER cholesterol: a delicate balance. Cell Metab. 8(6):512–521.

79. Duran JM, et al. (2012) Sphingomyelin organization is required for vesicle biogenesis at the Golgi complex. EMBO J. 31(24):4535–4546.

