## Supplementary Materials and Methods for "Reconstitution of ORP-mediated lipid exchange process coupled to PI(4)P metabolism"

#### 1 Building the kinetic model

In order to quantify the experimental observations described in the Results section, we established a kinetic model that describes in-vitro PI(4)P synthesis and hydrolysis as well as the transport of PI(4)P and DHE between ER and Golgi-like liposomes. To present the full kinetic model, we first introduce the kinetics for PI(4)P production (a) and hydrolysis (b) and for Osh4p-mediated lipid exchange (c).

##### 1.1 PI(4)P production

PI(4)P is synthesized in the membrane of Golgi-like liposomes through the phosphorylation of its precursor PI catalyzed by the PI4K construct (see the first section of Results). Given the abundance of PI in Golgi-like membranes, as well as ATP, largely in excess in buffer, we model this process as a first order reaction, and we ignore the depletion of PI (by the end of the experiment only a fraction of the initial concentration is phosphorylated into PI(4)P). The concentration of kinase sets the reaction rate ($\mathrm{k}_{kin}$):

$$\frac{dp_G}{dt} = \Pi$$

where $\Pi$ has units of $\mu M/s$ and increases when the concentration of kinase is larger. From experimental measures (Fig. 4B), we estimate that $\Pi \approx 7 \cdot 10^{-4} \mu M s^{-1}$ and $\Pi \approx 1.9 \cdot 10^{-3} \mu M s^{-1}$ for "low" and "high" kinase concentration respectively (i.e. $PIK/L = 1/200$ and $PIK/L = 1/100$). We assume that the kinase can phosphorylate PI multiple times, without undergoing degradation.

##### 1.2 PI(4)P hydrolysis

PI(4)P is hydrolyzed into PI by the protein Sac1[1-522]$_{\mathrm{His6}}$ attached to ER-like liposomes according to the following reaction:

$$\frac{dp_E}{dt} = -\mathrm{k}_{sac}\, p_E$$

where we assumed that water is abundant, that the concentration of Sac1 remains constant over time and that a single Sac1 copy can hydrolyze multiple molecules of PI(4)P. We estimate that $-\mathrm{k}_{sac} \approx 2.8 \times 10^{-3} s^{-1}$ for concentrations of Sac1$\approx 100\, nM$ (based on ref [1]).

##### 1.3 Lipid exchange on Osh4p

When Osh4p, in complex with one molecule of PI(4)P, encounters a membrane, it can drop its lipid cargo and pick up another one, thus changing the relative concentrations of complexes bound to PI(4)P ($p_o$) and to DHE ($s_o$) and of the lipids PI(4)P ($p$) and DHE ($s$) in this membrane:

$$op + s \rightleftharpoons os + p$$

If this exchange reaction is second order, the concentrations obey the following kinetics

$$\begin{cases} \dot{s} = -rp_o s + rs_o p \\ \dot{p} = rp_o s - rs_o p \\ \dot{s_o} = rp_o s - rs_o p \\ \dot{p_o} = -rp_o s + rs_o p \end{cases} \tag{1}$$

where $r$ is the rate at which Osh4p exchanges the lipids. The forward and backward reactions are assumed to have the same rate because Osh4p was estimated to have an equivalent affinity for DHE and PI(4)P [2], so it picks up these lipids based on their relative concentration. We designed a simple experiment to determine the rate ($r$). To this end, Osh4p-PI(4)P complex was mixed with liposomes containing different amounts of DHE. The substitution of PI(4)P by DHE in Osh4p, hence the exchange rate, was followed by measuring a FRET signal between tryptophan residues of the protein, surrounding the lipid-binding pocket, and the DHE molecule once loaded in this pocket [1, 2], (Supplementary Figure 4). Assuming that this reaction is at equilibrium, one single equation sums up the dynamics:

$$p_o s = s_o p \tag{2}$$

Further, Osh4p proteins and DHE are conserved, and PI(4)P is also conserved as no phosphorylation nor hydrolysis are allowed in this exchange essay, designed to measure $r$. Three additional equations express these conservation laws:

$$\begin{cases} s + s_o = S \\ p + p_o = P \\ s_o + p_o = O \end{cases} \tag{3}$$

where $S$, $P$ and $O$ are the total concentrations of sterol, PI(4)P and Osh4p in the system. Because PI(4)P is introduced in this exchange essay only through in complexed form with Osh4p, $O = P$. Solving the system by substitution with the initial condition $p(0) = 0$ leads to

$$s = \frac{S}{S+P}(S + P - O)\;,\; p = \frac{P}{S+P}(S + P - O) \tag{4}$$

and using that $P = O$ we obtain the equilibrium values for $s$ and $p$:

$$s = \frac{S^2}{S+O} \; , p = \frac{OS}{S+O} \tag{5}$$

Perturbing the equilibrium of this simplified system to the first order leads to the analytic solution for all concentrations, in particular for the concentration of Osh4p-sterol (i.e. Osh4p-DHE) complexes:

$$s_o(t) = \frac{OS}{S+O} - \frac{OS}{S+O} e^{-r(S+O)t} \tag{6}$$

Eq (6) describes well the experimental data, which confirms that the kinetics is accurately described by a second order reaction. Additionally, through the fitting procedure, we obtain an estimate for the rate of exchange $r = (1.5 \pm 0.1) \times 10^{-3}\ \mu M^{-1} s^{-1}$ (see Supplementary Figure 4).

### 2 Solving the kinetic model in vitro

To analyze the kinetics of lipid transfer between ER-like and Golgi-like liposomes, we develop a more complex model that hinges on the assumptions for production and hydrolysis of PI(4)P and lipid exchange described above. We define six unknown variables: the concentration of sterol in Golgi-like ($s_G$) and ER-like membranes ($s_E$); the concentration of PI(4)P in Golgi-like ($p_G$) and ER-like membranes ($p_E$); the concentration of Osh4p complexes loaded with sterol ($s_O$) and with PI(4)P ($p_O$) in the bulk. We assume that there are no free Osh4p proteins in our assays because the release of a lipid by Osh4p is instantaneously followed by the loading of Osh4p with another lipid molecule. In other words Osh4p is always bound to either sterol or PI(4)P.
Using the assumptions discussed above, the evolution of the concentrations obeys a set of ordinary differential equations:

$$\begin{cases} \dot{p}_E = -\mathrm{k}_{sac}\, p_E + p_o r s_E - s_o r p_E \\ \dot{s}_E = -p_o r s_E + s_o r p_E \\ \dot{p}_G = \Pi + p_o r s_G - s_o r p_G \\ \dot{s}_G = -p_o r s_G + s_o r p_G \\ \dot{p}_o = -p_o r s_E + s_o r p_E - p_o r s_G + s_o r p_G \\ \dot{s}_o = p_o r s_E - s_o r p_E + p_o r s_G - s_o r p_G \end{cases} \tag{7}$$

The first two equations of (7) show that $p_E$ increases (and $s_E$ decreases) if a complex $p_o$ from the bulk hits a ER-like membrane, binds to DHE and releases its PI(4)P cargo into the membrane. Similarly, $p_E$ decreases (and $s_E$ increases) in the opposite case, where a $s_o$ complex releases its DHE cargo to bind to PI(4)P. On a ER-like membrane the concentration of PI(4)P decreases in time because of the process of hydrolysis. The third and fourth equation in the system (7) are similar to the first two equations but relate to kinetics on Golgi-like liposomes except that PI(4)P is synthesized in the Golgi-like membrane

whereas it is degraded in ER-like membranes.
By summing respectively, the last two equations and the second fourth and sixth equation we realize that there are two conserved quantities in the system:

$$\begin{cases} s_o + p_o = O \\ s_E + s_G + s_o = S \end{cases} \tag{8}$$

where $O$ and $S$ are the total concentration of sterol and proteins respectively and are fixed within each experiment.

##### Solution in vitro, short times

The full model can be solved asymptotically or integrated numerically. We introduce two non-dimensional parameters $A = k_{sac}/rS$ and $B = \Pi/rS^2$ and rewrite the system of equations in a non-dimensional formulation

$$\begin{cases} \dot{p}_E = -Ap_E + p_o s_E - s_o p_E \\ \dot{p}_G = B + p_o s_G - s_o p_G \\ \dot{s}_E = -p_o s_E + s_o p_E \\ \dot{s}_G = -p_o s_G + s_o p_G \\ \dot{p_o} = s_o(p_G + p_E) - p_o(s_E + s_G) \\ \dot{s_o} = -s_o(p_G + p_E) + p_o(s_E + s_G) \end{cases} \tag{9}$$

Note that from now on, all concentrations are normalized to $S$ and times are normalized with $(rS)^{-1}$. In the interest of clarity, we do not introduce a different symbol for the non-dimensional variables which would would make the notation considerably heavier. Unless otherwise stated, we consider from now on all variables nondimensionalized. In our experiments, at time 0 no PI(4)P is present and as time proceeds it is produced in Golgi-like membranes ($L_G$). Therefore we can not assume $\dot{p}_G$ to be at the equilibrium. As soon as $p_G$ is produced in $L_G$ membranes, the exchange starts and, since the lipid transfer assays are done in stirring conditions, we assume that the exchange reaction (last two equations in system (9)) is at equilibrium i.e. $p_o s_E - s_o p_E + p_o s_G - s_o p_G = 0$. From the fourth equation in system (9):

$$p_o = (\frac{O}{S} - p_o)\frac{p_G + p_E}{1 - O/S - p_o} \sim \frac{O}{S}\frac{p_G + p_E}{1 - O/S} \tag{10}$$

To get to equation (10) we took advantage of the conservation laws for both DHE and Osh4p $s_E + s_G + s_o = 1$, and $s_o + p_o = O/S$, and we approximated $p_o \sim 0$ which is valid at short times. The time derivative of $p_o$ reads:

$$\dot{p}_o = \frac{O}{S - O}(\dot{p}_G + \dot{p}_E) = \frac{O}{S - O}(B - A\frac{p_o s_E}{A + O/S - p_0}) \tag{11}$$

where we considered $p_E$ at the equilibrium, hence from the first equation in system (9) $p_E = p_o s_E/(A+s_o)$. The combining equilibrium of the first equation with the third equation in system (9) we obtain:

$$\dot{s}_E = -Ap_E = -A\frac{p_o s_E}{A + O/S - p_o} \tag{12}$$

We next neglect $p_0 \sim 0$ at the denominator in eqs (11) (12) (short times) and introducing three non-dimensional parameters $\eta = B\frac{O}{S}$, $\beta = \frac{OA}{SA+O}$, $\gamma = \frac{A}{A+O/S}$ we obtain:

$$\begin{cases} \dot{p}_o = \frac{O}{S}(B - A\frac{p_o s_E}{A+s_o}) = \eta - \beta p_o s_E \\ \dot{s}_E = -A\frac{p_o s_E}{A+s_o} = -\gamma p_o s_E \end{cases} \tag{13}$$

$$\begin{cases} p_o = x \\ s_E = \frac{1}{2} + y \end{cases} \tag{14}$$

By separation of variables:

$$\frac{dx}{\eta - \frac{1}{2}\beta x} = dt \rightarrow x(t) = \left(x_0 - \frac{\eta}{\frac{1}{2}\beta}\right) e^{-\frac{1}{2}\beta t} = \frac{2\eta}{\beta}(1 - e^{-\frac{1}{2}\beta t})$$

$$dy = -\frac{1}{2}\frac{2\eta\gamma}{\beta}(1 - e^{-\frac{1}{2}\beta t})dt \rightarrow y = y_0 - \frac{\eta\gamma t}{\beta} + x\frac{\gamma}{\beta}$$

In dimensional units the solutions reads:

$$s_E(0) - s_E(t) = \Pi\left(t - 2\tau * (1 - e^{-t/2\tau})\right) \tag{15}$$

where $\tau = \frac{k_{sac}+Or}{rOk_{sac}}$. At short times, this equation implies that sterol exchange is not equal to PI(4)P production, $\Pi t$, but rather it is decreased by a quantity $2\Pi\tau(1 - e^{-t/(2\tau)})$. All other variables can now be obtained by substitution. In Figure Supplementary 5, we show that the asymptotic solution is in perfect agreement with the numerics at short times and yields a value for the timescale $\tau = 684 \pm 9s$ . Note that because this is a short time solution, $\tau$ is independent on the value of $\Pi$. In Figure Supplementary 6, we show the evolution of the system at longer times, in a regime that is discussed next.

##### Solution in vitro, long times

At long times, the system of equations (9) predicts that all sterol is transferred from ER- to Golgi-like membranes. However, we noted that our experiments show a non-trivial long time behavior with a finite fraction of sterol in ER-like membranes. In order to account for this system, we included the spontaneous transfer of sterol between ER- and Golgi-like membranes due to the direct exchange occurring upon collision of the liposomes in the stirring buffer. The full system accounting for spontaneous transfer is:

$$\begin{cases} \dot{p}_E = -\mathrm{k}_{sac}\, p_E + p_o r s_E - s_o r p_E \\ \dot{s}_E = -p_o r s_E + s_o r p_E \\ \dot{p}_G = \Pi + p_o r s_G - s_o r p_G - r_s(s_G - s_E) \\ \dot{s}_G = -p_o r s_G + s_o r p_G + r_s(s_G - s_E) \\ \dot{p}_o = -p_o r s_E + s_o r p_E - p_o r s_G + s_o r p_G \\ \dot{s}_o = p_o r s_E - s_o r p_E + p_o r s_G - s_o r p_G \end{cases} \tag{16}$$

where $r_s$ is the rate of spontaneous transfer. At long times, the system reaches a non-trivial equilibrium where:

$$\frac{S}{2}\left(1 - \frac{O}{S} - \frac{\Pi}{Sr_s}\right) \leq s_E \leq \frac{S}{2}\left(1 - \frac{\Pi}{Sr_s}\right)$$

where we used that $S - O \leq s_E + s_G \leq S$ and $s_G - s_E = \Pi/(Sr_s)$. To obtain an estimate of the rate of exchange, we fix $\Pi \approx 7 \times 10^{-4}$ and $1.9 \times 10^{-3} \mu M/s$, which matches the final production of $2.6\,\mu M$ and $6.7\,\mu M$ PI(4)P after 1 hour, for small and high kinase concentration respectively. Using $S = 10\,\mu M$ and $O = 0.2\,\mu M$, and $s_E^{\infty} = 4.2\,\mu M$ and $3.5\,\mu M$ from our experiments for small and high kinase concentration respectively, we obtain:

$$\frac{\Pi}{S - 2s_E^{\infty}} \leq r_s \leq \frac{\Pi}{S - O - 2s_E^{\infty}}$$

$$(4.4 \times 10^{-4}) \leq r_s \leq (4.9 \times 10^{-4})s^{-1} \quad \text{low kinase concentration}$$

$$(6.3 \times 10^{-4}) \leq r_s \leq (6.8 \times 10^{-4})s^{-1} \quad \text{high kinase concentration}$$

### 3 Model *in vivo*

In the cell, we can assume that sterol dynamics at the ER/Golgi interface is at equilibrium, due to sterol synthesis in the ER membrane and export from the trans-Golgi by vesicular trafficking. The system of equations (7) used to analyze the kinetics in vitro cannot be used directly to describe this system because sterol would accumulate indefinitely in the Golgi membranes as a function of PI(4)P synthesis. Similar to what was done for spontaneous transfer, it was necessary to introduce two additional terms: a constant production rate of sterol in the ER membrane, E, and a constant rate of export of sterol from the Golgi apparatus, D:

$$\begin{cases} \dot{p}_E = -Ap_E + p_o s_E - s_o p_E \\ \dot{p}_G = B + p_o s_G - s_o p_G \\ \dot{s}_E = -p_o s_E + s_o p_E + E \\ \dot{s}_G = -p_o s_G + s_o p_G - D \\ \dot{p_o} = s_o(p_G + p_E) - p_o(s_E + s_G) \\ \dot{s_o} = -s_o(p_G + p_E) + p_o(s_E + s_G) \end{cases} \tag{17}$$

We define the exchange rate at the ER as $p_o s_E - s_o p_E = t_E$ and similarly the exchange rate at the Golgi as $p_o s_G - s_o p_G = t_G$. When gradients need to first be established, PI(4)P production is needed to promote the formation of Osh4p-PI(4)P complexes (or OSBP-PI(4)P complexes in the case of human cells), and initiate the dynamics, as discussed for the model in vitro. When the cell reaches equilibrium, PI(4)P production is only needed to offset the transport of sterol from the trans-Golgi which amounts to a loss of sterol in the Golgi. In other words, if there was no export from the Golgi apparatus ($D = 0$), the dynamics would remain at equilibrium with no need for further transport nor PI(4)P production and consumption.

Let us now assume that there is an established gradient of sterol (fix $s_E$ and $s_G = 1 - s_E$) and let us assume there is a nonzero export of sterol from the trans-Golgi, i.e. $D > 0$: in these conditions, sterol needs to be transported to the Golgi apparatus, and this requires PI(4)P production. Mathematically, at equilibrium $B = -t_G = t_E = E = -D = Ap_E$ which means that to maintain the set sterol gradient, PI(4)P and sterol production, PI(4)P hydrolysis and lipid transport between ER and Golgi membrane must match (remember that the non-dimensional parameters are defined as $A = k_{sac}/rS$ and $B = \Pi/rS^2$). More precisely: when one sterol molecule is exported away from the trans-Golgi, one sterol molecule must be produced at the ER and transported from the ER to the Golgi membrane; this requires that one molecule of PI(4)P is produced in the Golgi membrane, transported from the Golgi to the ER membrane and then hydrolyzed by Sac1 on the ER membrane.

Next we can ask: what is the maximum sterol export from the Golgi apparatus that can be offset by increasing transport of sterol from the ER to the Golgi? To answer this question we solve for the state of equilibrium. For a given sterol gradient, i.e. given $s_E$ and $s_G = 1 - s_E$, few lines of algebra yield:

$$p_o = \frac{B}{S}\frac{AS + O}{B + s_E A} \tag{18}$$

$$s_o = \frac{O}{S} - p_o \tag{19}$$

$$p_E = \frac{B}{A} \tag{20}$$

$$p_G = \frac{p_o s_G + B}{s_o} \tag{21}$$

First, we show that given the concentration of protein, there is a constraint on the maximum value of $\Pi$. To clarify the constraint, let us consider the following limits. When $B \to 0$, $p_o \to 0$, $p_E \to 0$, $s_o \to O/S$ and $p_G \to 0$ i.e. if the system lacks PI(4)P no Osh4p-PI(4)P (or OSBP-PI(4)P) complexes are formed and no transport occurs, hence maintenance of equilibrium is only possible if no depletion of sterol on Golgi occurs. In the opposite limit, when $B \to \infty$, from equation (18) we have that $p_o \to A + O/S > O/S$, but this value of $p_o$ is larger than the (nondimensional) concentration of proteins $O/S$. Thus in order to maintain this equilibrium, more Osh4p-PI(4)P or OSBP-PI(4)P complexes are needed than there are available. This shows that there is a limit to the amount of leakage of sterol from the Golgi apparatus that can be replenished by transport from the ER. The maximum concentration of Osh4p-PI(4)P or OSBP-PI(4)P complexes is equal to the total concentration of proteins:

$$p_o^* = \frac{O}{S} = B^* \frac{AS + O}{S(B^* + s_E A)} \rightarrow B^* = s_E \frac{O}{S} = \frac{\Pi^*}{rS^2} \rightarrow \Pi^* = rO\bar{s}_E \tag{22}$$

where we used the overbar to denote the dimensional concentration of sterol on the ER, $\bar{s}_E$, in $\mu M$ (note that in the main text, $\bar{s}_E$ is indicated with $s_E$, as all concentrations are dimensional and we do not use the overbar). $\Pi^*$ is the maximum production rate of PI(4)P, equal to the maximum export of sterol from the Golgi apparatus that can be replenished by transport of sterol from the ER. In plain words eq (22) means that sterol depleted from Golgi membranes can be replenished by transport from the ER only up to the rate $rO\bar{s}_E$. More sterol can be transported by speeding up transport (increasing $r$); increasing the number of transport proteins ($O$) or of sterol in the ER membrane ($\bar{s}_E$).

In the case of Hela cells and yeast, we have estimates for $s_G$, $p_G$, $O$, $Sac1$. Because $p_E$ is typically very small, and from equation (20), $p_E = B/A$, we assume that $B/A << 1$, hence we work in the limit $B << A$. Under this limit, plugging equations (18) and (19) into equation (21) we obtain that:

$$p_G \approx \frac{ABS + BOs_G}{Os_E A - ABS}$$

hence

$$B \approx \frac{Os_E A p_G}{AS + Os_G + ASp_G}$$

and remembering that $B = \Pi/(rS^2)$ and $A = k_{sac1}/(rS)$ we obtain:

$$\Pi = \Pi^* \frac{p_G k_{sac1}}{p_G k_{sac1} + rOs_G + k_{sac1}} \tag{23}$$

From equation (23) note that the actual rate of PI(4)P production, $\Pi$, is always smaller than $\Pi^*$ because the fraction that multiplies $\Pi^*$ on the right-hand side of equation (23) is clearly smaller than 1. Note that the assumption $B << A$ is verified for Hela and yeast cells ($B = 6 \times 10^{-6}$ and $A = 10^{-4}$ for yeast, and $B = 1.3 \times 10^{-5}$ and $A = 1.6 \times 10^{-4}$ for the first estimate on Hela cells, and $B = 1.8 \times 10^{-5}$ and $A = 6 \times 10^{-5}$ for the second estimate on Hela cells).

### References


[1] de Saint-Jean, M, Delfosse, V., Douguet, D., Chicanne, G., Payrastre, B., Bourguet, W., Antonny, B., Drin, G. "Osh4p exchanges sterols for phosphatidylinositol 4-phosphate between lipid bilayers" *J Cell Biol* **195**: 965–978 (2011).

[2] Moser von Filseck, J., Vanni, S., Mesmin, B., Antonny, B., Drin, G. "A phosphatidylinositol-4-phosphate powered exchange mechanism to create a lipid gradient between membranes" *Nat. Comm.* **6**:6671 (2015)
